# *BioISO*: an objective-oriented application for assisting the curation of genome-scale metabolic models

**DOI:** 10.1101/2021.03.07.434259

**Authors:** Fernando Cruz, João Capela, Eugénio C. Ferreira, Miguel Rocha, Oscar Dias

## Abstract

As the reconstruction of Genome-Scale Metabolic Models becomes standard practice in systems biology, the number of organisms having at least one metabolic model at the genome-scale is peaking at an unprecedented scale. The automation of several laborious tasks, such as gap-finding and gap-filling, allowed to develop GSMMs for poorly described organisms. However, such models’ quality can be compromised by the automation of several steps, which may lead to erroneous phenotype simulations.

The Biological networks constraint-based *In Silico* Optimization (*BioISO*) is a computational tool aimed at accelerating the reconstruction of Genome-Scale Metabolic Models. This tool facilitates the manual curation steps by reducing the large search spaces often met when debugging *in silico* biological models. *BioISO* uses a recursive relation-like algorithm and Flux Balance Analysis to evaluate and guide debugging of *in silico* phenotype simulations. The potential of *BioISO* to guide the debugging of model reconstructions was showcased using GSMMs available in literature and compared with the results of two other state-of-the-art gap-filling tools (*Meneco* and *fastGapFill*). Furthermore, *BioISO* was used as *Meneco*’s gap-finding algorithm to reduce the number of proposed solutions (reaction sets) for filling the gaps.

*BioISO* was implemented as a webserver available at https://bioiso.bio.di.uminho.pt; and integrated into *merlin* as a plugin. *BioISO’s* implementation as a Python™ package can also be retrieved from https://github.com/BioSystemsUM/BioISO.

## Background

The reconstruction of Genome-Scale Metabolic Models (GSMMs) is becoming a standard practice in systems biology. GSMMs can be used to simulate the organism’s phenotype under different environmental and genetic conditions [1–3]. Flux Balance Analysis (FBA) [4], or related constraint-based methods, are used for solving linear programming problems outlined by constraints imposed over the stoichiometric model.

Nevertheless, reconstructing GSMMs is still challenging [1], as model validation and manual curation can be laborious tasks [3]. Most bottlenecks derive from accumulated errors, which require complex and unique solutions. For instance, when a metabolic network is converted into a stoichiometric model, FBA often mispredicts the organism’s experimental growth rate due to errors like missing or blocked reactions and dead-end metabolites (gaps), among others.

The reconstruction of GSMMs can follow two diverse paradigms: bottom-up [1] and top-down [5]. The widely-adapted bottom-up paradigm consists of four main steps: draft reconstruction based on genome functional annotation; refinement and curation of the draft reconstruction; conversion to stoichiometric model; model validation [1]. The last steps of a bottom-up reconstruction usually include several time-consuming and repetitive tasks aimed at fixing errors that emerged during the draft reconstruction, so that the discrepancy between the predicted phenotype and experimental results can be solved. While on one hand errors can be solved using manual curation, there are several gap-find and gap-fill tools to accelerate the debugging process.

Most state-of-the-art tools for debugging draft reconstructions comprehend both automated gap-finding and gap-filling procedures [6–9]. Nevertheless, there are other tools developed for only one of these procedures [10–12]. Besides, gap-find and gap-fill tools can also be separated according to the gap-finding and gap-filling methodology. Whereas gap-find algorithms aim to find either missing or blocked reactions and dead-end metabolites in a draft reconstruction, gap-fill tools are responsible for finding potential solutions to the errors mentioned above.

Regarding gap-finding methodologies, *biomassPrecursorCheck* [10], and *Meneco* [6] are based on guided-search algorithms to identify gaps or errors directly associated with a given objective/reaction. Both tools check the metabolic network topological features to assert gaps. That is, these tools assert the existence of predecessors and successors of a given metabolite. The *COBRA Toolbox*’s tool *BiomassPrecursorCheck* searches for predecessors immediately downstream of the biomass reaction of a given model, whereas *Meneco* accelerates the gap-search according to a set of seed and target metabolites provided as input. However, the search depth of the latter algorithm may encompass the whole metabolic network.

*gapFind*/*gapFill* [7], *fastGapFill* [8], and *Gauge* [9] are based on exhaustive searches. Thus, these methodologies identify gaps all over the metabolic network, regardless of a given objective. *gapFind*/*gapFill* and *fastGapfill* highlight gaps using a stoichiometry-like approach. These methods search the stoichiometric matrix for no-production and no-consumption metabolites. Alternatively, Gauge combines Flux Coupling Analysis and gene expression data to propose gaps in a draft GSMM.

To the best of our knowledge, all gap-gill tools require a dataset of metabolic reactions, usually retrieved from a biochemical database (e.g. KEGG [13], BiGG [14] or MetaCyc [15]), to fulfil metabolic gaps [6–9, 11, 12]. Besides a database of metabolic reactions, both *Gauge* [9] and *Mirage* [12] require gene expression data. *Smiley* [11] relies on additional growth phenotype data to identify minimal environmental conditions for which the model mispredicted growth and non-growth phenotypes. *Meneco*, *gapFind/gapFill*, *fastGapFill*, *Gauge* and *Smiley* consider the minimal reaction set of the whole dataset to fulfil each single gap. Alternatively, *Mirage* considers a pan-metabolic network that assures flux through all metabolites but then applying a pruning step to reduce the large set of solutions. Thus, the solution set is often the result of two very different gap-filling approaches, namely the parsimonious and pruning approaches.

Most state-of-the-art tools for debugging draft reconstructions rely on proprietary software, such as MATLAB (Mathworks®) or GAMS. *Meneco* is the only tool freely available to the community, as it is available as a Python package. It is worth noticing though that all tools require coding skills to be used. More importantly, the main output of these tools consists of excessively verbose outputs such as large arrays of missing metabolites and even greater sets of potential solutions.

Most gap-fill tools warn that gaps might result from missing mappings between the metabolites’ abbreviations and the reference database identifiers. Besides the mapping’s limitations, several tools require different format-files for the metabolic data, such as SBML (e.g., Meneco), KEGG reaction database *lst* format file (e.g., fastGapFill), customised text files (e.g., gapFind/gapFill) or data structures (e.g., Gauge). On the other hand, other tools lack information on how a different source of solutions can be used (e.g., Mirage and Smiley).

Besides the lack of easy-to-use computational tools, most tools often require long runtimes and return highly verbose, complex, and extensive outputs. The examination of these results can be challenging for wet-lab scientists who do not have coding skills nor data analysis expertise.

Several gap-find and gap-fill state-of-the-art tools have been described with further detail in the Supplementary Files S1-2.

Unlike the bottom-up paradigm, the fast and automated top-down paradigm does not resort to gap-filling procedures. This alternative approach consists of reconstructing an universal GSMM that has been curated previously for most common errors [5]. This universal simulation-ready model is then converted to an organism-specific model by carving reactions and metabolites for which evidence is missing. Thus, the top-down paradigm can be extremely useful to create microbial community models by merging the automated single-species models into community-scale networks [5]. Nevertheless, it is still not clear whether single-species models’ phenotype simulations are unreasonably biased by the universal GSMM.

The introduction of artefacts in metabolic networks can hinder GSMM’s applications, such as metabolic engineering and drug targeting tasks. These issues may be extremely relevant for organisms that have evolved due to a combination of extensive loss-of-function events and acquisition of key genes, via horizontal gene transfer during co-evolution with well-defined and constant ecological niches [16–18]. Moreover, although loss-of-function genetic variants are frequently associated with severe clinical phenotypes, several events are also present in healthy individuals’ genomes, making it essential to assess their impact [19].

Hence, automated approaches, and especially gap-fill tools, must be used very carefully according to the constraints raised herein. Otherwise, the offered automation can be a counterproductive solution for the manual curation steps performed during high-quality reconstructions. Furthermore, the usability of gap-fill approaches can be vastly improved.

To the best of our knowledge, the reconstruction of high-quality GSMMs is often based on a parsimonious bottom-up approach, involving manual curation and human intervention. In our view of a parsimonious bottom-up reconstruction, the metabolic network can be divided into smaller, yet insightful, modules based on the phenotype being studied. Then, recursive relations can be used to accomplish the division of metabolic networks into the smaller modules directly associated with the objective phenotype. FBA simulations applied over surrogate reactions designed explicitly for each module can unveil the minor manual curation tasks that often increment the quality of the reconstruction and resolve the metabolic gaps for such module.

With this methodology in mind, *BioISO* has been designed to automatize the search for reactions and metabolites associated with a given objective, narrowing the search space. These reactions and metabolites are properly evaluated by *BioISO* that uses FBA over surrogate reactions. Then, the results are presented into a user-friendly manner, so that the debugging process can be easy to follow and repeat.

## Implementation

### *BioISO*’s algorithm

*BioISO* requires a constraint-based model and the reaction to be evaluated, which defines the objective function of the linear programming problem to be solved. A recursive relation-like algorithm is then used to build a hierarchical structure according to the metabolites and reactions associated with this objective.

*BioISO* is herein showcased through the analysis of a small-scale metabolic network represented in Figure 1, and having 12 intracellular and two extracellular metabolites, 12 reactions, and two compartments (extracellular and intracellular). In this metabolic network, the reaction identified by *R_8_* is considered missing, blocked, or incorrectly formulated, while identifier *R_11_* refers to the reaction to be evaluated.

**Figure 1.**
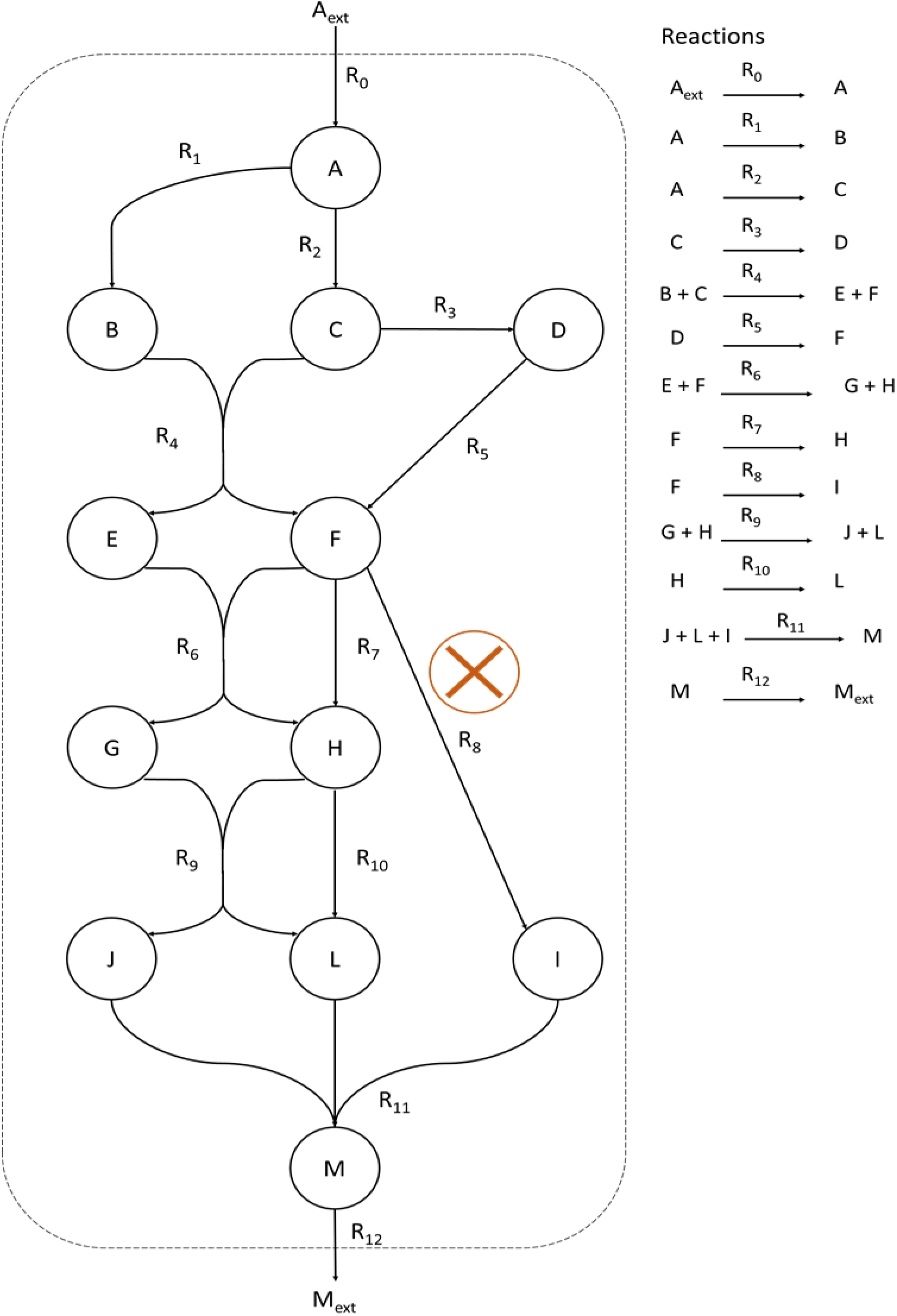
Small-scale metabolic network. Metabolites and reactions are represented in the metabolic network as white nodes and black directed arrows. The extracellular boundary is represented as a dashed line. The reactions are listed alongside the metabolic network. In this metabolic network, the reaction identified by *R_8_* is considered missing, blocked, or incorrectly formulated.

*BioISO* starts by finding the set of metabolites associated with the reaction submitted for evaluation, namely the reaction *R_11_*. The tool will find metabolites *J*, *L, I* and *M* as next nodes since these metabolites are involved in *R_11_* (Figure 2). A set of reactions is then created for each node and populated with the reactions associated with each metabolite. Thus, *BioISO* will retrieve four sets of reactions, one for each metabolite (Figure 2).

**Figure 2.**
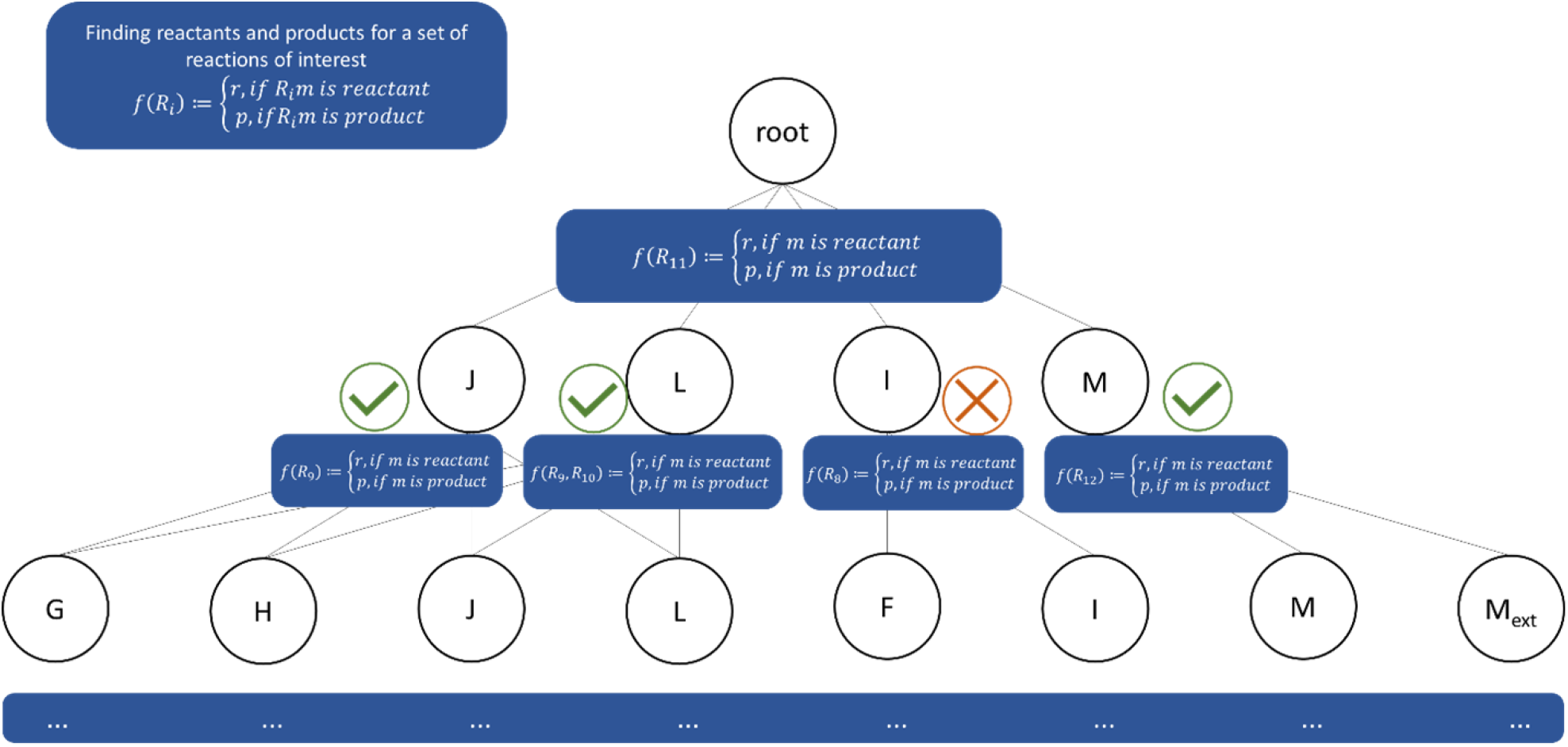
Evaluation of reaction *R_11_* with *BioISO*. *BioISO* finds the set of metabolites associated with reaction *R_11_*, which in this case corresponds to metabolites *J*, *L, I,* and *M*. For the next call, *BioISO* finds metabolites *G*, *H*, *J, L, F*, *I, M,* and *M_ext_* in the reactions *R_8_*, *R_9_*, *R_10_* and *R_12_* and so forth.

Meanwhile, *BioISO* assembles a hierarchical tree-based structure, depicted in Figure 3. The tool identifies as precursors (reactants) or successors (products) the metabolites associated with the submitted reaction (R11). Thus, *J*, *L,* and *I* (reactants) and *M* (product) involved in rection *R_11_* were separated into two different branches: precursors and successors, respectively.

**Figure 3.**
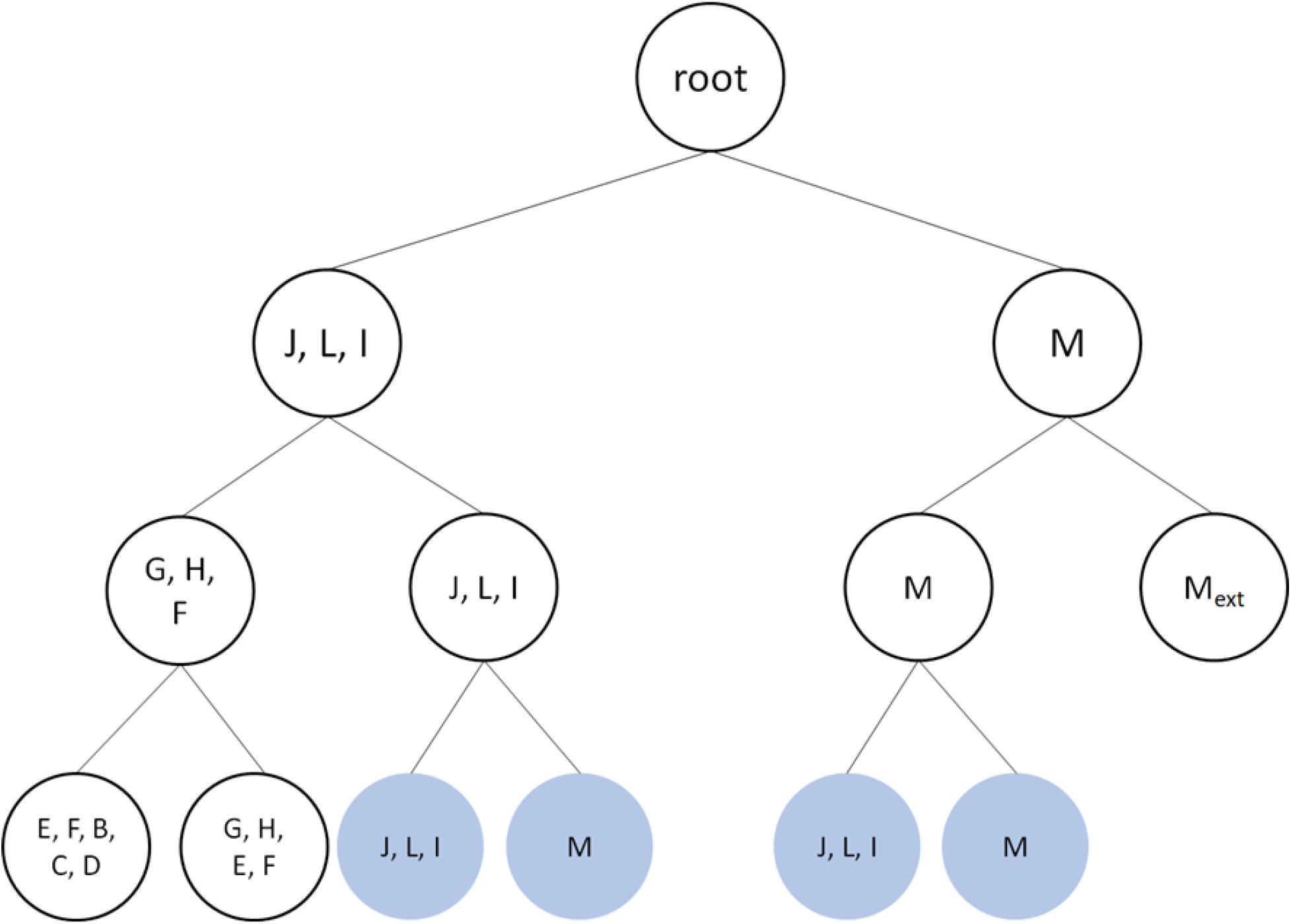
The hierarchical tree-based structure, imposed by the recursive relation-like computational method implemented in *BioISO*, is outlined. *BioISO* finds the set of precursors and successors in the first level, which in this case correspond to metabolites *J*, *L, I,* and *M*, respectively. For the next level, *BioISO* finds the precursors *G*, *H*, *F* for the previous precursors *J*, *L, I*, which are also successors of themselves. On the other branches, *M* is its own precursor, while *M_ext_* is the successor. *BioISO* has implemented a cache memory system of all simulations performed during the recursion. Thus, nodes coloured in blue are only evaluated once, as they were already evaluated in those specific conditions.

In the next recursive call, *BioISO* retrieves metabolites *G*, *H*, *F, M*, *J*, *L*, *I,* and *M_ext_* from reactions *R_8_*, *R_9_*, *R_10_*, and *R_12_* (Figure 2), while adding the precursors *G*, *H*, *F,* and *M*, and the successors *J*, *L*, *I,* and *M_ext_* to the tree-based structure (Figure 3). These reactions are either consuming or producing the metabolites identified in the previous step (*J*, *L, I* and *M*).

The stopping condition, namely *BioISO’s* depth, represents the number of recursive calls performed during the analysis of the metabolic network. For instance, varying *BioISO*’s depth from 1 to 3 allows running the tool for shallow, guided or nearly exhaustive searches, depending on the metabolic network’s size and arborescence.

The methodology for finding and assessing metabolites and reactions is detailed in algorithms S3.1-4 of the Supplementary File S3. The first algorithm labelled *BioISO* (algorithm S3.1) is the core logic supporting the methodology proposed in this work. *BioISO* uses algorithm S3.2 to find and evaluate (using FBA) reactions associated to nodes. More importantly, *BioISO* uses a more comprehensive approach to evaluate reactants (precursors) and products (successors), as demonstrated in algorithms S3.3 (*testReactant*) and S3.4 (*testProduct*), respectively.

In detail, a **precursor (reactant)** is considered as a positive assessment if the metabolic model can **produce** it (connected metabolite). Thus, a reactant is a product elsewhere in the metabolic network; otherwise, it would not be available for the objective reaction. Hence, *BioISO* evaluates an unbalanced reaction explicitly designed to allow the metabolite accumulation in the metabolic model. The evaluation is successful if the model can attain a positive flux in the FBA solution for this surrogate reaction.

For example, when evaluating reaction *R_11_*, *BioISO* will evaluate the precursor *I* by adding an unbalanced reaction *R_13_*, which takes *I* as a reactant, and whose lower and upper bounds are set to zero and plus-infinity, respectively.

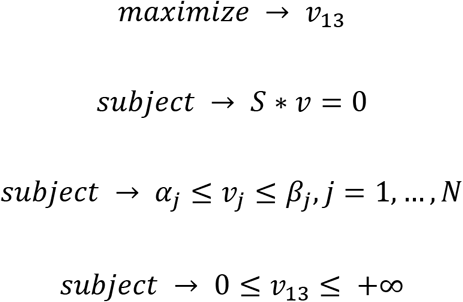

Where:

- *v_13_* is the linear objective function for maximisation of reaction *R_13_*;
- *v* is the flux vector;
- *S* is the stoichiometric matrix (columns represent reaction fluxes and rows the metabolites mass balances);
- α and β are the lower and upper bounds, respectively.

Furthermore, a similar reaction is included in the model for each reactant, to prevent the seldom cases in which all reactants are forcibly produced by reactions that produce/synthesise the assessed metabolite.

Likewise, *BioISO* creates unbalanced reactions that allow the uptake of all products associated with the evaluated reaction. These reactions are included in the model to cover up for the unlikely scenario that the model forcibly needs to consume/metabolise such products to synthesise the precursor.

On the other hand, a **successor (product)** is considered as a positive assessment if the metabolic model can **consume** it (connected metabolite). Thus, a product is a reactant elsewhere in the metabolic network. As described in the testing of precursor *I*, *BioISO* also creates an unbalanced reaction for the successor. However, this reaction is now explicitly designed to allow the metabolite **uptake** in the metabolic model. Thus, the minimisation of this uptake reaction is now the objective function of the FBA simulation. In other words, the model should now metabolise/consume the precursor metabolite, obtaining an optimal non-zero flux solution through the unbalanced reaction.

A detailed description of *BioISO* workflow to search and assert gaps is provided in the Supplementary File S3.

In short, the procedure to split the objective into two sub-problems and evaluate both metabolites and reactions follows the workflow below:

1. collect the reactions associated with each metabolite to be evaluated;
2. maximise/minimise the reactions and assess the outcome of the FBA solution;
3. from such reactions, find the precursor (reactants) and successor (products) metabolites;
4. create unbalanced reactions allowing accumulation or uptake of the metabolites;
5. maximise/minimize the unbalanced reactions and assess the outcome of the FBA solutions.

An analysis of BioISO’s relation-like algorithm’s complexity, together with the recursion tree method visualisation, is also provided in Supplementary File S3.

### *BioISO*’s applications

*BioISO* is a package developed in Python™ 3 using the FBA framework implemented in COBRApy [20]. *BioISO* relies on COBRApy to read GSMMs written in the System Biology Markup Language (SBML) [21]. The IBM CPLEX solver (v. 1210) is used by default to solve multiple linear programming problems formulated with the FBA framework, although any solver supported by COBRApy can be used. *BioISO*’s source code, validation procedures and examples can be obtained from our group’s GitHub at https://github.com/BioSystemsUM/BioISO.

A Dockerised Flask application has been implemented to make *BioISO* available to all scientific community at https://bioiso.bio.di.uminho.pt. This webservice allows one to submit a GSMM in the SBML format-file to be evaluated for a specific reaction available in the model. Then, *BioISO*’s webservice will return a user-friendly webpage highlighting the metabolic network’s blocked reactions and dead-end metabolites according to the submitted reaction. Finally, the user is encouraged to navigate through the set of dead-end metabolites in an intuitive manner.

Besides the webservice application, *BioISO* is also available as a plugin for *merlin* [22]. This plugin allows using *BioISO* to supply an equally user-friendly view of the errors associated with a given model being reconstructed within *merlin*.

Finally, Supplementary File S4 provides instructions to run *BioISO* in the available applications and to interpret the expected results.

## Results

*BioISO* is aimed at identifying errors that emerge during the bottom-up reconstruction of high-quality GSMMs. Errors such as missing or blocked reactions and dead-end metabolites are often met during model debugging and refinement. Thus, *BioISO* is based on a recursive-like algorithm to guide the search for metabolic gaps associated with a given objective. Throughout *BioISO*’s objective oriented search, multiple FBA simulations are used to assert real metabolic gaps. Hence, we purpose a tool capable of reducing large search spaces and asserting real metabolic gaps to accelerate time-consuming and laborious manual curation tasks.

Most state-of-the-art tools for debugging draft reconstructions aim to find and solve a wide range of problems. These tools are commonly used in automatic gap-find and gap-filling routines. For instance, *Meneco*, *gapFind/gapFill*, *fastGapFill*, *Gauge*, *Smiley* and *Mirage* are gap-fill tools aimed at finding and solving errors accumulated during the draft reconstruction.

*gapFind/gapFill*, *fastGapFill* and *Gauge* exhaustive-search tools attempt to assert gaps throughout the whole metabolic network. Then, these tools enumerate minimal solutions (set of reactions) to solve the highlighted gaps. Alternatively, Mirage and Smiley add new reactions to the model without an initial gap-scan, forcing model predictions to match the experimental data.

On the other hand, Meneco guide-search algorithm searches for gaps according to a set of seed and target metabolites. Then, this tool enumerates a minimal set of reactions that can restore the flux to all dead-end metabolites identified during the topological search.

Likewise, *BioISO* seeks dead-end metabolites downstream and upstream of a user-defined objective. Additionally, *BioISO* performs multiple FBA’ simulations of custom unbalanced reactions during the topological search to evaluate whether a given metabolite is actually being consumed or produced.

More importantly, *BioISO* is the only tool available to all scientists. That is, *BioISO* is the only user-friendly gap-finding tool, providing a graphical user interface embedded in both a webserver and *merlin*. Thus, our tool allows users to analyse gaps and errors without requiring coding skills or additional metabolic data such as growth phenotype data or biochemical databases. Moreover, BioISO is a ready-to-use and relatively fast method, allowing users to run this tool iteratively during model reconstruction.

A summary of all features used to compare *BioISO* with several gap-find and gap-filling tools is available in Supplementary Files S1-2.

Furthermore, *BioISO’s* validation includes three assessments: *BioISO’s* algorithm depth analysis; Exhaustive-search *versus* guided-search; BioMeneco – embedding *BioISO* in Meneco [6]. The first analysis was aimed at assessing BioISO’s shallow, guided, or nearly exhaustive searches for metabolic gaps in five state-of-the-art models. The second analysis allowed to assess the relevance of guided- and exhaustive-searches for gap-finding. In this assessment, we have compared *BioISO* and *Meneco* guide-searches against *fastGapFill* exhaustive-search. Finally, the last analysis showcases the outcome of setting *BioISO* as Meneco’s gap-finding algorithm.

### BioISO’s algorithm depth analysis

*BioISO* was used to analyse several published GSMMs for two objective functions: growth and compound production maximisation. The sets of dead-end metabolites and blocked reactions were determined as described in the Implementation section. The workflow and methodology used to assess *BioISO*’s algorithm robustness is described in further detail in the Materials and Methods section together with Supplementary Files S5-6.

*BioISO’s* analysis included different settings, namely varying the algorithm’s depth from 1 to 3 allowing to set *BioISO* for shallow, guided or nearly exhaustive searches.

*BioISO’s* depth level of 1 assesses the nearest neighbours (successors and precursors metabolites) and their associated reactions. According to Figure 4, *BioISO* analyses less than 50% of all reactions for growth maximisation. The number of blocked reactions is significantly reduced at depth level 1 (less than 12%), except for the iJO1366 [23] and iBsu1103 [24] models (Figure 4 and Supplementary Files S6.1-2). Likewise, *BioISO* only covers less than 10% of all metabolites for growth maximisation (Figure 4). As a result, the number of dead-end metabolites found by *BioISO* at a depth level of 1 is less than 3 for all models (Figure 4 and Supplementary Files S6.1-2). The level of insight provided by *BioISO* for shallow searches is significantly reduced and similar to the biomassPrecursorsCheck tool from COBRA Toolbox [10] or Meneco [6].

**Figure 4.**
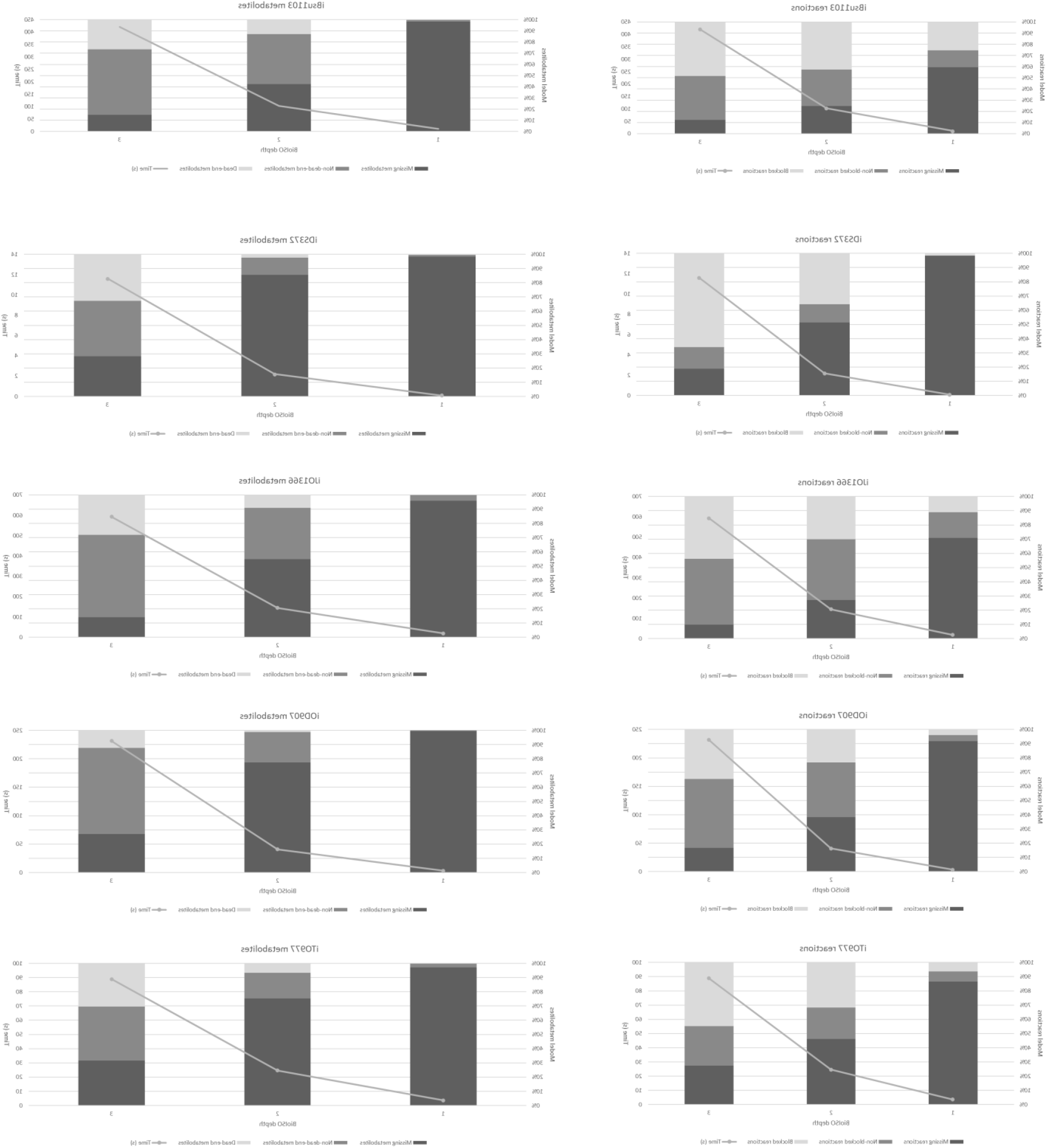
Summary of the reactions (left panel) and metabolites (right panel) analysed by *BioISO* for published Genome-Scale Metabolic Models. *BioISO* was used to analyse 5 state-of-the-art models (iBsu1103, iDS372, iJO1366, iOD907 and iTO977) with added gaps. *BioISO’s* analysis included different algorithm settings, namely varying the depth from 1 to 3 for each objective function, which will control the number of recursive calls for precursors and successors. This allowed to run *BioISO’s* algorithm for shallow, guided or nearly exhaustive searches, depending on the size and arborescence of the metabolic network. *BioISO*’s computation time was recorded in seconds (s) together with missing (non-covered by *BioISO*) reactions and metabolites, non- and dead-end metabolites, non- and blocked reactions.

Increasing the depth level to 2 allows evaluating more reactions. As demonstrated in Figure 4, 50% or more of the reactions are assessed in all models. Whereas *BioISO* analyses nearly a quarter of all metabolites in the iOD907 [24] and iTO977 [25] models, this coverage increases up to 60% in the iJO1366 [25] and iBsu1103 [24] models. In contrast, only 15% of all metabolites have been covered by *BioISO* in the iDS372 [26] model. *BioISO* also detects more blocked reactions and dead-end metabolites for guided searches. The percentage of blocked reactions varies between 20% to 40%, and the percentage of dead-end metabolites between 2% to 12% (Figure 4 and Supplementary Files S6.1-2).

At a depth level of 3, a nearly exhaustive search is performed as *BioISO* analyses more than 70% of both metabolites and reactions (Figure 4 and Supplementary Files S6.1-2). Likewise, the percentage of detected blocked reactions and dead-end metabolites increases up to 65% and 30%, respectively.

As detailed in the Supplementary Files S6.3-4, similar results were obtained for the maximisation of compound production. However, the number of metabolites covered in the iTO977 model is considerably lower than the remaining models at depth levels of 2 and 3 (Supplementary File S6.4).

Figure 4 also presents *BioISO’s* computation time for each model during growth maximisation, as a function of the depth level. *BioISO* was considerably faster for shallow searches (depth level of 1) in all models for both growth (Figure 4 and Supplementary Files S6.1-2) and compound production (Supplementary Files S6.3-4) maximisation. According to Figure 4, *BioISO* required computation time for a depth level of 2 varies between 2 to 144 seconds during growth maximisation. Whereas, during the compound production maximisation, *BioISO* takes between 4 to 74 seconds (Supplementary Files S6.3-4). The computation time of *BioISO* increases significantly at the depth level of 3. At this depth, *BioISO’s* computation time can attain around 600 and 405 seconds when maximising growth (Figure 4 and Supplementary Files S6.1-2) and compound production (Supplementary Files S6.3-4), respectively.

Hence, *BioISO’s* computation time is significantly dependent on the size of the covered search space. In turn, the covered search space increases with the depth of search and the model’s size. Although navigating the network throughout the new metabolites and reactions might not be time-consuming, evaluating numerous metabolites and reactions using the FBA framework requires time.

The dead-end metabolites and blocked reactions ratios were calculated as denoted in equations 1), 2), 3) and 4) of the Materials and Methods section. Figure 5 highlights the ratios of dead-end metabolites obtained for both growth and compound production analysis in all models. Both dead-end metabolites and blocked reactions ratios are also available at the Supplementary Files S6.1-4.

**Figure 5.**
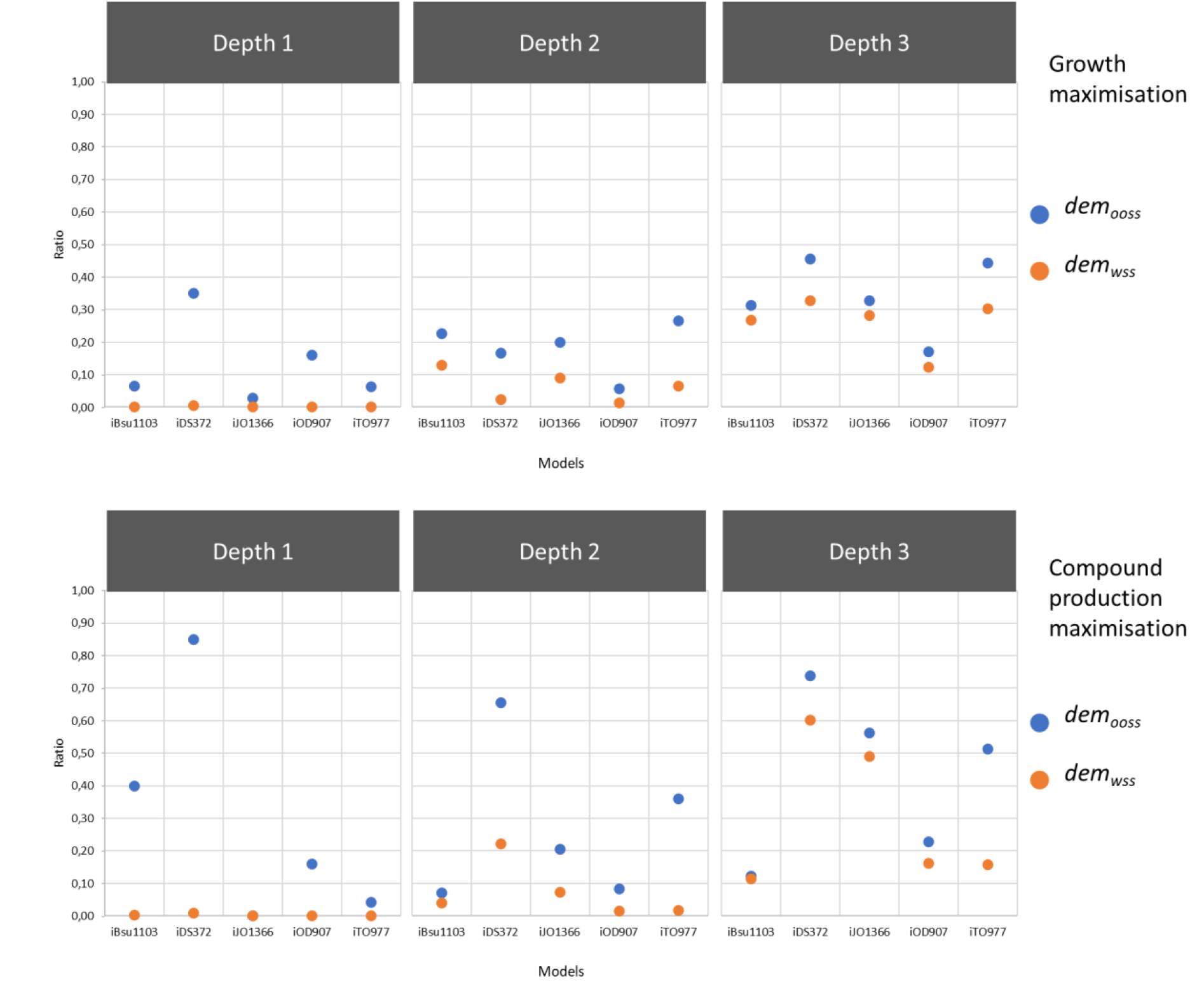
Calculated ratios of dead-end metabolites for the maximisation of growth (upper panel) and compound production (bottom panel). *BioISO* was used to analyse 5 state-of-the-art models (iBsu1103, iDS372, iJO1366, iOD907 and iTO977) with added gaps. *BioISO’s* analysis included different algorithm settings, namely varying the depth from 1 to 3 for each objective function, which will control the number of recursive calls for precursors and successors. This allowed to run *BioISO’s* algorithm for shallow, guided or nearly exhaustive searches, depending on the size and arborescence of the metabolic network. *dem_ooss_* and *dem_wss_* stand for the ratios of dead-end metabolites for the objective-oriented search and whole search spaces.

At a depth level of 1, the ratio of blocked reactions for the objective-oriented search (*br_oos_*) varies between 0.3 and 0.9, whereas the homologous ratio for the whole-space search (*br_wss_*) varies between 0.02 and 0.25 (Supplementary Files S6.1-4).

Regarding the ratios of dead-end metabolites, *BioISO* attains markedly small ratios for the whole-space search *(dem_wss_*) at a depth level of 1, namely obtaining ratios smaller than 0.1 in all models for both objective functions (Figure 5 and Supplementary Files S6.1-4). However, the ratio of dead-end metabolites for the objective-oriented search *(dem_oos_*) can peak up to 0.35 (Figure 5 and Supplementary Files S6.1-2) and 0.85 (Figure 5 and Supplementary Files S6.6-4) in the growth and compound production analysis, respectively.

The *br_wss_* ratios increase significantly when raising the depth to level 2 (*BioISO* guided search), whereas *br_oos_* ones remain roughly the same as in the previous level. The *br_wss_* ratio can vary from 0.23 to 0.43 (Supplementary Files S6.1-2) and from 0.23 to 0.36 (Supplementary Files S6.3-4) for the growth and compound production analysis, respectively.

The *dem_oos_* ratio tends to increase as a response to *BioISO’*s guided search (depth level of 2) in the iBsu1103, iJO1366, and iTO977 models during the growth maximisation analysis (Figure 5). In contrast to the previous trend, the *dem_oos_* ratio tends to decrease in the iDS372 and iOD907 models. Regarding the maximisation of compound production, *BioISO* also attains higher *dem_oos_* ratios in both iJO1366 and iTO977 models at a depth level of 2 (guided search). Nevertheless, the *dem_oos_* ratio obtained in the iBsu1103 model is smaller in comparison to the value obtained for the shallow search (depth level of 1).

In general, the *dem_wss_* ratio tends to increase as a response to *BioISO’*s guided search (depth level of 2) in all models for both objective functions, though not exceeding 0.221. As shown in Figure 5 and Supplementary Files S6.1-2, the *dem_wss_* ratio ranges between 0.01 (iOD907 model) and 0.13 (iBsu1103 model) for the growth maximisation analysis. During the compound production maximisation analysis, the *dem_wss_* ratio is less than 0.1 in all models except for the iDS372 model where it peaks 0.221 (Figure 5 and Supplementary File S6.3-4).

Using *BioISO* for nearly exhaustive searches (depth level of 3) returns *br_oos_* ratios between 0.41 and 0.81, whereas the *br_wss_* ratio varies between 0.35 and 0.66 for both objective functions (Supplementary File S6.1-4). As for detecting dead-end metabolites during the growth maximisation analysis, *BioISO* nearly exhaustive search attains the highest *dem_oos_* and *dem_wss_* ratios of 0.455 and 0.327 in the iDS372 model (Figure 5), respectively. Regarding the compound production maximisation, *BioISO* peaked for a depth level of 3 *dem_ooss_* and *dem_wss_* ratios of 0.737 and 0.602 in the iDS372 model (Figure 5), respectively.

Although the *br_oos_* ratio oscillates when rising depth, the *br_wss_* tends to increase steadily. Similarly, the *dem_wss_* also tends to increase with depth for both objective functions, whereas the *dem_oos_* ratio mimics the oscillatory behaviour of the *br_oos_* ratio. The oscillatory behaviour of the *br_oos_* and *dem_oos_* ratios is heavily pronounced between depth 1 and 2, which can be associated with the reduced level of detail that *BioISO* can provide for shallow searches.

The high *br_wss_* ratios obtained for all levels are likely associated with the fact that *BioISO* does not prevent circular dependencies nor by-products accumulation when testing reactions. The interactive output of *BioISO* in both webserver and *merlin* guides the user through the dead-end metabolites (precursors and successors having an unsuccessful evaluation) while providing the evaluation of the reactions for guidance and further insight.

When testing metabolites, *BioISO’s* strategy to prevent circular dependencies and by-product accumulation as well as the isolated evaluation of precursors and successors seems to have a greater impact on reducing the set of dead-end metabolites. The *dem_wss_* ratio is significantly smaller for shallow and guided searchers across all models for both objective functions.

Furthermore, although *BioISO* has attained *dem_wss_* ratios higher than 0.4 for two models during the compound production maximisation analysis with a depth level of 3, this ratio remains below 0.33 in all models during the growth maximisation analysis. The *dem_wss_* ratios higher than 0.4 obtained with nearly exhaustive searches of *BioISO* may be associated with the factual metabolic gaps that do not need to be corrected or might not be associated with the desired phenotype.

For example, *BioISO* has systematically attained higher ratios, for all metrics, when assessing the iDS372 incomplete models for both objective functions. These higher scores may be associated with poor connectivity of most metabolites involved in the metabolic pathways analysed by *BioISO*, as parasitic organisms evolve in rich media, thus developing auxotrophies [16, 26, 27]. Hence, it is worth noticing that the identified dead-end metabolites might be associated with real metabolic gaps that should not be gap-filled.

In short, *BioISO* scores most of the smaller *dem_wss_* ratios at the depth level of 2 (guided search). Moreover, the gap between *dem_wss_* and *dem_oos_* ratios also starts to narrow for *BioISO*’s guided search. The small difference between both metrics suggests that most dead-end metabolites suggested by *BioISO* are a direct outcome of the actual network gaps introduced during the validation. Hence, *BioISO* can suggest a higher number of dead-end metabolites associated with the objective, while maintaining the curation efforts at a minimum.

Therefore, a depth level of 2 was selected as the default level for running *BioISO*, after analysing all *dem_wss_*. Using this depth value, *BioISO* can guide the search to identify errors in a given metabolic network, without evaluating only the direct precursors and successors, such as *biomassPrecursorsCheck* [10] and Meneco [6], or the burden of evaluating the whole network, such as fastGapFill [8] and gapFind/gapFill [7].

Furthermore, a significant part of the metabolic network associated with a given objective is analysed by the tool for a guided search (depth level of 2), while the required computation time is significantly lower.

### Exhaustive-search versus guided-search

The exhaustive-search versus guided-search assessment was designed to compare the results of two guided-search tools, namely *BioISO* (guided search – depth level of 2) and Meneco [6], with fastGapFill [8] exhaustive-search application. For that, the iJO1366 and iDS372 models obtained for the growth maximisation analysis were used in this assessment. The workflow and methodology used to compare exhaustive-searches against guided-searches is described in further detail in the Materials and Methods section together with Supplementary Files S5-6. Figure 6 exhibits a summary of dead-end metabolites and blocked reactions ratios calculated for each tool.

**Figure 6.**
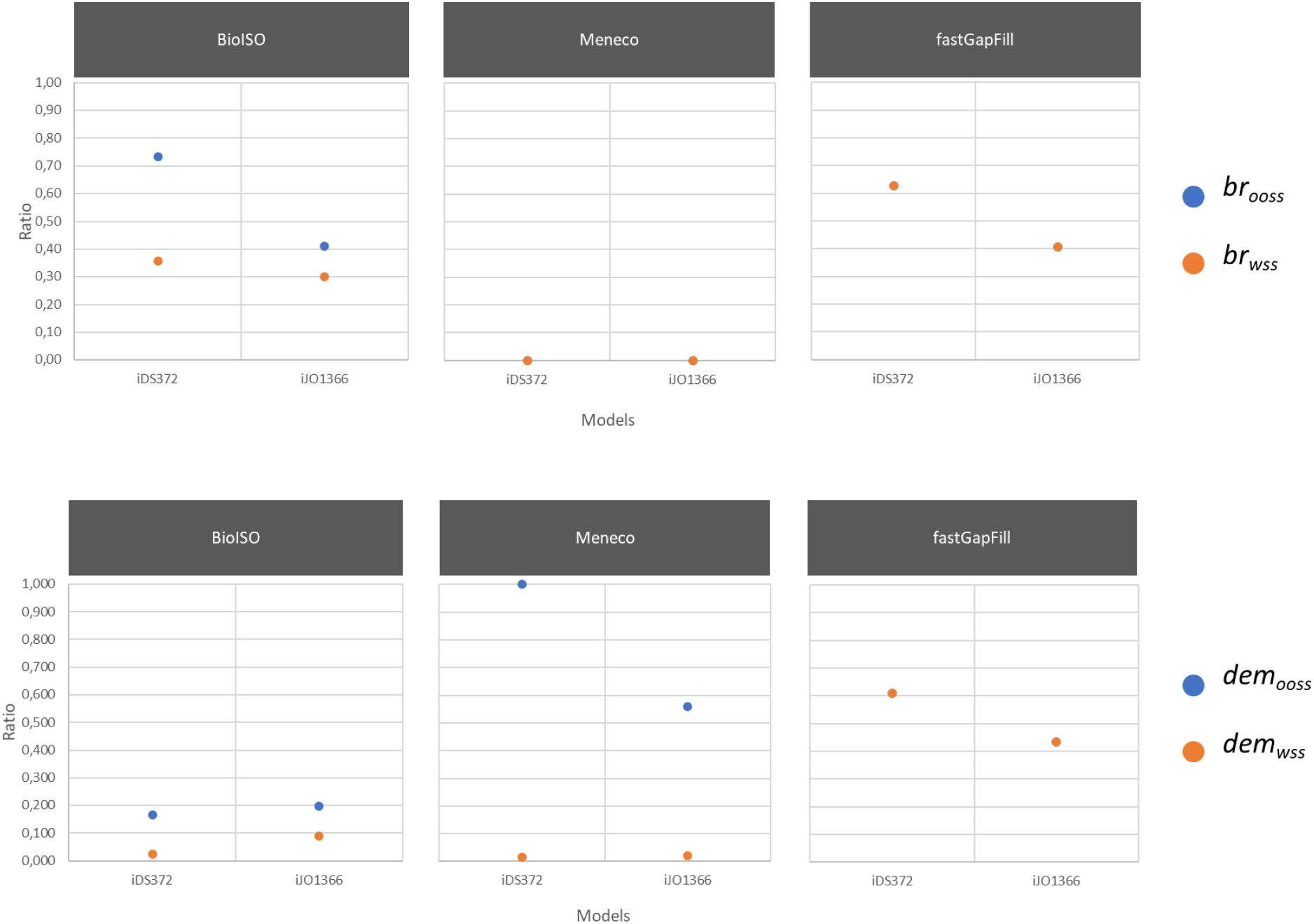
Assessment of the relevance of guided (*BioISO* and Meneco) versus exhaustive searches (fastGapFill) for gap-finding. *BioISO*, Meneco and fastGapFill were used to highlight gaps in two state-of-the-art models (iDS372 and iJO1366), with added gaps. The ratios of dead-end metabolites (*dem_ooss_* and *dem_wss_*) and blocked reactions (*br_ooss_* and *br_wss_*) for both objective-oriented search (*ooss*) and whole search (*wss*) spaces were then calculated for each tool.

According to Figure 6 and Supplementary Files S6.5-9, Meneco performed the poorest in identifying metabolic gaps. Besides the poor performance regarding the assessment of dead-end metabolites, Meneco does not provide insights on blocked reactions.

The number of covered metabolites when using Meneco is the same as the number of metabolites selected for target metabolites, as this tool only evaluates target metabolites. No information is provided about other metabolites in the metabolic network.

Hence, although Meneco obtained the lowest *dem_wss_* ratios in both models (Figure 6), the tool suggests the absence of biosynthetic pathways to synthesise all metabolites in the covered search space, thus obtaining the highest *dem_oos_* ratios in both models (Figure 6). In short, most metabolites analysed by this tool were evaluated as dead-end metabolites in the incomplete models.

fastGapFill provides, on the other hand, a level of insight much larger than Meneco, analysing all reactions and metabolites in all models (Figure 6 and Supplementary Files S6.5-9). However, the exhaustive-search tool is associated with two significant drawbacks. Firstly, fastGapFill is the slowest tool (Supplementary Files S6.5-9). Secondly, this gap-filling tool highlights numerous blocked reactions and dead-end metabolites according to Figure 6, which might hinder a fast and precise identification of a *de facto* error in the network, such as the ones introduced in this validation procedure.

For example, fastGapFill has attained higher *dem_wss_* ratios (Figure 6) for the model of the less described and smaller genome’s organism (*Streptococcus pneumoniae*), which has probably evolved through a combination of extensive loss-of-function events during the co-evolution with well-defined and constant ecological niches [16, 26, 27].

Although Meneco has obtained smaller *dem_wss_* scores than *BioISO*, the level of insight provided by the gap-filling tool for both metabolic networks is significantly lower than the detail provided by our tool. When comparing the *dem_oos_* ratios, it is clear the lack of insight provided by Meneco, as most of the metabolites covered by Meneco are highlighted as dead-end metabolites. On the other hand, *BioISO* can be more effective and precise by suggesting fewer dead-ends out of the examined metabolites pool (Figure 6 and Supplementary Files S6.5-9).

According to Figure 6, *BioISO* attained lower *br_wss_* and *dem_wss_* ratios than fastGapFill in both models. Hence, such smaller *br_wss_* and *dem_wss_* ratios suggest that *BioISO* is more capable of reducing the whole-space search to fewer dead-end metabolites than the gap-filling tool.

When debugging and validating the model for specific objective functions, such as growth maximisation, *BioISO* seems better suited for reducing the search space for errors and gaps in metabolic networks, than the other tools analysed in this assessment. This advantage allows spending less time debugging unrealistic errors or gaps. Furthermore, as *BioISO* reduces the search space for errors, it also favours parsimonious alterations to the draft metabolic network. As a result, *BioISO* can be of paramount importance for the high-quality bottom-up reconstruction of GSMMs during the manual curation stage.

However, it should be noticed that Meneco and fastGapFill have been designed to be essentially gap-fill tools. Thus, these tools use different approaches than *BioISO* to find errors.

### BioMeneco – embedding BioISO in Meneco

*BioMeneco*, *BioISO’s* integration with Meneco [6], was developed to determine whether the former can improve the latter results by narrowing the search space for the gap-filling task. For that, reactions “R04568_C3_cytop” and “SO4tex”, were removed from the iDS372 and iJO1366 models, respectively. Meneco was then used to generate potential solutions for restoring models’ prediction of a growth phenotype based on *BioISO* suggestions for the set of targets (primary input for Meneco).

All metabolites identified as not being produced or consumed by *BioISO* in the iDS372 model are reported in Table 1. These metabolites have been selected for the set of target metabolites after a brief analysis of the *BioISO’s* output.

**Table 1.** Metabolites identified by *BioISO* as not being produced or consumed (dead-end metabolites) by the iDS372 model missing R04568_C3_cytop reaction.

| Dead-end metabolite | Connected dead-end metabolites |
| --- | --- |
| C01356_cytop (Lipid) | C05980_cytop; C06040_cytop; C04046_cytop;<br>C00344_cytop |
| C06042_cytop (Lipoteichoic acid) | C00116_cytop; C00162_cytop |

In the iDS372 model, *BioISO* indicated that reaction “R04568_C3_cytop” might be associated with the synthesis of a precursor of the lipidic and lipoteichoic acid pathways. Most dead-end metabolites identified by *BioISO* were associated with metabolites “C01356_cytop” and “C06042_cytop”, which are biomass precursors representing the lipid and lipoteichoic acid cellular biomass fractions, respectively. These suggestions are in agreement with the metabolites being synthesised by the reaction removed from the model. Reaction “R04568_C3_cytop” is associated with the synthesis of “trans-Tetradec-2-enoyl-[acp]” (“C05760_cytop”), which in turn is a precursor of the compound “Tetradecanoyl-[acp]” (“C05761_cytop”) involved in the fatty acid biosynthesis pathway.

Besides the dead-end metabolites shown in Table 1, *BioISO* suggested an unsuccessful evaluation of all remaining biomass precursors and successors. Nevertheless, only the lipid and lipoteichoic acid compounds were identified as dead-end metabolites. The remaining biomass precursors and successors refer to the special cases reported in Supplementary File S4. Briefly, these metabolites were unsuccessfully evaluated due to a missing or impaired reaction downstream, namely the biomass reaction.

The corresponding precursors and successors of all neighbour metabolites were classified as non-dead-end metabolites, except for several precursors and successors of the lipid and lipoteichoic acid compounds.

All metabolites identified as not being produced or consumed by *BioISO* in the incomplete iJO1366 model, reported in Table 2, were selected for the set of target metabolites.

**Table 2.**
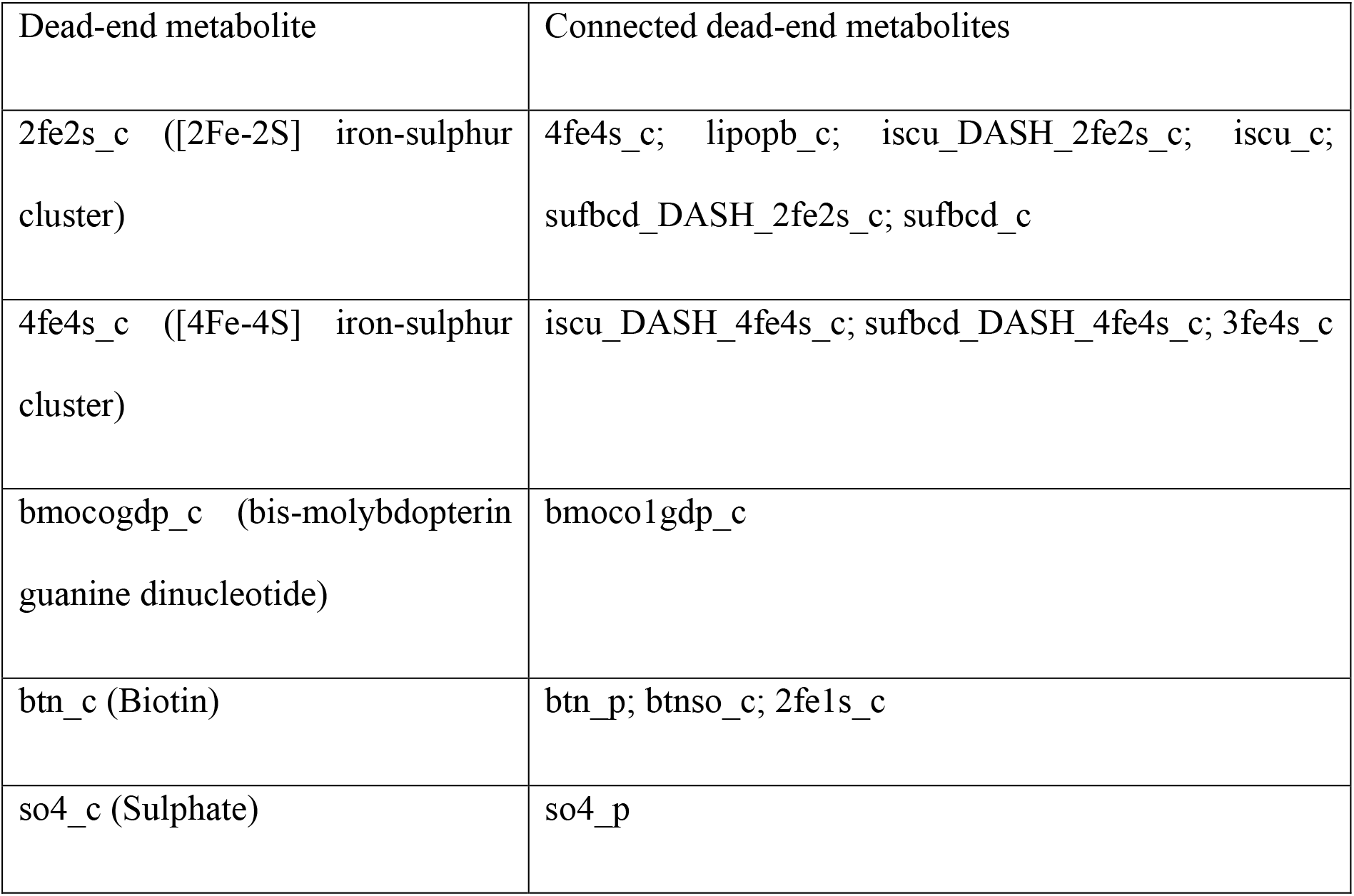
Metabolites identified by *BioISO* as not being produced or consumed (dead-end metabolites) by the iJO1366 model missing SO4tex reaction.

*BioISO* has indicated that the missing “SO4tex” reaction might be associated with the synthesis of a precursor for the sulphur metabolism. Most dead-end metabolites identified by *BioISO* were linked to the iron-sulphur clusters, biotin, bis-molybdopterin guanine dinucleotide, and sulphate biomass precursors, which are all associated with the sulphur requirements of *E. coli*. These results are in line with the transport of sulphate to the periplasm, by the removed reaction. The “SO4tex” reaction is responsible for transporting sulphate from the extracellular medium to the periplasm, which is then transported to the cytoplasm.

*BioISO* suggested more dead-end metabolites than the precursors described in Table 2, which are absent from the set of “target” metabolites for the iJO1366 model. *BioISO* negatively evaluated the biomass precursors “mobd_c”, “sheme_c”, “cl_c”, and “2ohph_c”. Nevertheless, these metabolites have been ignored as dead-end metabolites, as they refer to the special cases reported in Supplementary File S4. In fact, the precursors of these metabolites are being produced and the successors being consumed, except for the biomass. Thus, *BioISO* highlighted these metabolites because the biomass reaction is, actually, the only consumption site available. These metabolites are an example of unsuccessful evaluations that should be easily detected in the user-friendly output returned by the webserver and *merlin*. Moreover, the automatic tools would deal with such cases as perhaps regular gaps and try to incorrectly resolve them.

The gap-filling solutions suggested by Meneco for the incomplete iDS372 model are satisfactory, as the proposed solutions could restore flux through the biomass reaction, and thus through all set of targets initially proposed. Meneco suggests adding reaction “R04568” (which was previously removed for this test) to restore the metabolic model.

Nevertheless, other solutions may add artefacts in the iDS372 model. Reactions “R11633”, “R09085”, “R11636”, “R11634”, “R11671” and “R00183” are equally recommended to restore flux throughout the set of targets. However, these reactions are not involved in the synthesis or consumption of missing biomass precursors or successors. Most reactions are involved in the synthesis of biomass precursors not affected by the removed reaction, such as the “R11636” (“dCTP” synthesis), “R09085” (carbon metabolism), “R11634” (“dATP “synthesis), and “R11633” (“dGTP” synthesis). Other reactions are involved in the synthesis of metabolites not required for *S. pneumoniae’s* growth. Note that, only reactions suggested in the pool named “One minimal completion” were considered. Nevertheless, Meneco provides a complete enumeration of all combinations of minimal completions.

*BioMeneco* recommended, on the other hand, a reduced pool of gap-filling solutions. In this case, *BioMeneco* suggested reaction “R04568”, but now only three reactions (“R11671”, “R00182”, and “R09085”) have been equally proposed. As neither RNA nor DNA were included in the set of target metabolites, all reactions previously suggested to restore the synthesis of purines and pyrimidines have been discarded.

Meneco restored the test iJO1366 model for six biomass precursors while indicating 35 “unreconstructable targets”. Meneco identified the “so4_c” metabolite, which is one of the biomass precursors affected by the removal of the “SO4tex” reaction, as “reconstructable”. Nevertheless, the remaining metabolites for which Meneco could restore flux were not affected by the removed reaction.

More importantly, Meneco’s output does not comprise the “SO4tex” reaction in the set of gap-filling solutions to restore flux through all biomass precursors. More surprisingly, the “SO4tex” reaction was not included in any combination of minimal completions obtained through the complete enumeration of solutions. Furthermore, other potential solutions can lead to the introduction of artefacts in the iJO1366 model. For example, all reactions included in the “One minimal completion” set of solutions are transport reactions for biomass precursors not affected when reducing the iJO1366 model.

Interestingly, only the reaction “SO4tex” has been suggested by *BioMeneco* to restore flux through all missing biomass precursors and successors in the test iJO1366 model. As this time none of the other biomass precursors was included in the set of target metabolites, all reactions involved in the transport of co-factors, ions, and amino acids were discarded from the “One minimal completion” pool.

Therefore, *BioISO* can be used to decrease large search spaces associated with model debugging procedures. Besides proposing a user-friendly application to guide the search for dead-end metabolites, we have showcased that *BioISO* can also facilitate high-quality bottom-up reconstructions by adjusting the guided-search gap-filling tool Meneco. For that, we suggest BioMeneco as an iterative process comprising two separate tasks:

1. running *BioISO* to identify the set of metabolites not being produced or consumed (dead-end metabolites).
2. running Meneco using the set of metabolites highlighted earlier as target metabolites, to obtain parsimonious solutions to complete draft metabolic networks.

## Conclusions

*BioISO* is a user-friendly tool, capable of performing guided searches of gaps in metabolic networks. This tool aims to assist the reconstruction of high-quality genome-scale metabolic models by scientists without coding skills, leveraging bottom-up reconstructions that require manual curation and human intervention.

Several state-of-the-art gap-find and gap-fill tools have been surveyed with *BioISO*, which emerged as the only open-source tool ready to be used by everyone in the scientific community. Moreover, *BioISO* is not associated with the significant drawbacks of using a gap-filling method, such as poor usability, requirement for additional data, and recommending biological artefacts due to the lack of evidence for the solutions.

*BioISO* has been validated with GSMMs available in the literature [23, 24, 26, 28, 29]. *BioISO’s* validation comprehended three assessments: algorithm depth analysis – assessment of shallow, guided or nearly exhaustive searches with *BioISO*; exhaustive-search (fastGapFill [8]) versus guided-search (*BioISO* and Meneco [6]); embedding *BioISO* in Meneco [6] - the outcome of setting *BioISO* as Meneco’s gap-finding algorithm.

The ratio of dead-end metabolites in the whole-space search increases with the depth of search. Nevertheless, *BioISO* attains lower ratios for shallow (depth level of 1) and guided searches (depth level of 2). Moreover, a significant part of the metabolic network associated with a given objective is analysed by the tool at a depth level of 2, which provides the best trade-off.

Although *BioISO* has attained lower dead-end metabolites ratios in the whole-space search than fastGapFill, Meneco has scored even smaller ratios. Nevertheless, the level of detail provided by Meneco is significantly lower, as most of the metabolites covered by Meneco are highlighted as dead-end metabolites. On the other hand, *BioISO* can be more effective and precise by suggesting fewer dead-ends from the examined metabolites pool.

When debugging and validating the model for specific objective functions, such as growth maximisation, *BioISO* seems better suited for reducing the search space for errors and gaps in metabolic networks, than the other tools analysed in this assessment.

When using *BioISO* to pre-process Meneco’s set of targets, the latter suggested the correct minimal completions. *BioISO* can improve Meneco’s gap-finding algorithm, facilitating Meneco’s integration with a high-quality bottom-up reconstruction workflow by following an iterative process comprising two separate tasks: running *BioISO* to identify dead-end metabolites; running Meneco using the set of metabolites highlighted as target metabolites, to obtain parsimonious solutions.

## Materials and Methods

### BioISO’s algorithm depth analysis

The analysis of *BioISO’*s algorithm depth was performed in parallel for both objective functions, namely growth and compound production maximisation. This assessment allowed setting shallow, guided or nearly exhaustive searches with *BioISO* to assess the algorithm’s robustness. *BioISO’*s algorithm depth analysis was performed in five state-of-the-art GSMMs: iDS372 (*Streptococcus pneumoniae*) [26]; iJO1366 (*Escherichia coli*) [23]; iBsu1103 (*Bacillus subtilis*) [24]; iTO977 (*Saccharomyces cerevisiae*) [29]; iOD907 (*Kluyveromyces lactis*) [28].

For instance, five incomplete models were created for the growth maximisation analysis by removing the following reactions from the *E. coli* iJO1366 model [23], one at a time: SDPTA; IMPC; MEPCT; NNDPR; SERAT. The incomplete models were evaluated by setting *BioISO*’s algorithm depth level at 1, and growth maximisation as the objective function (Ec_biomass_iJO1366_core_53p95M). This procedure was repeated for the remaining models and the depth levels of 2 and 3. Similarly, this procedure was repeated for the compound production maximisation analysis.

Then, two metrics were proposed to evaluate the gap-finding performance, namely the ratio of dead-end metabolites and the ratio of blocked reactions. These metrics were used to quantify the search space associated with model debugging of gaps and errors. As shown in equation 1), the ratio of dead-end metabolites for the objective-oriented search space (*dem_ooss_*) is a function of the number of metabolites that a guided-search tool evaluates as unsuccessful (dead-end metabolite) divided by the size of the objective-oriented search space (*ooss*). The ratio of dead-end metabolites for the whole search space (*dem_wss_*) is a function of the number of found dead-end metabolites divided by the *wss*, as described in equation 2). Equations 3) and 4) describe a similar approach to calculate the ratio of blocked reactions for the objective-oriented search (*br_ooss_*) and the whole search (*br_wss_*) spaces, respectively.

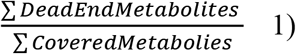

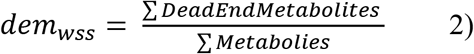

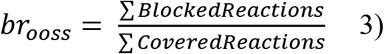

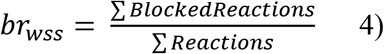

Objective functions, settings, evaluation metrics and methodologies used to introduce gaps in state-of-the-art GSMMs used during the algorithm depth analysis can also be consulted in detail at Supplementary Files S5-6.

### Exhaustive-search versus guided-search

The exhaustive-search versus guided-search analysis was performed for both iDS372 [26] and iJO1366 [23] models to assess the relevance of guided (*BioISO* and *Meneco* [6]) and exhaustive searches (*fastGapFill* [8]) for gap-finding.

*BioISO* was used as described in the previous section for a depth level of 2.

All metabolites available in the extracellular compartment of the iDS372 or iJO1366 models were used as seed metabolites to run *Meneco* gap-finding methodology. Likewise, the set of target metabolites was comprised of precursors and successors of the evaluated reactions. The set of dead-end metabolites was determined through *Meneco’s* ‘*get_unproducible*’ method. Note that, *Meneco* cannot assert blocked reactions. Thus, the set of blocked reactions could not be determined.

*fastGapFill* was used to assess the whole search space by accounting for errors and gaps. *fastGapFills’ ‘gapFind’* and *‘findBlockedReaction’* methods were used to determine dead-end metabolites and blocked reactions, respectively, in the incomplete models.

The metrics described in the previous section were then used to assess the tools’ performance, namely the ratio of dead-end metabolites and the ratio of blocked reactions. Supplementary Files S5-6 present all details about the assessment of *BioISO* with *Meneco* and *fastGapFill*.

### BioMeneco – embedding BioISO in Meneco

The BioMeneco analysis was performed for the iDS372 [26] and iJO1366 [23] models to assess the integration of *BioISO* as *Meneco’s* gap-finding algorithm.

*Meneco’s* topological search finds dead-end metabolites, so the gaps associated with them can be filled with reactions from a universal database. The novelty of *Meneco* is that it allows selecting which gaps should be filled by tweaking the set of target metabolites. Hence, *BioMeneco*, *BioISO’s* integration with *Meneco*, was performed to assess whether *BioISO* can suggest the right set of targets to be used as input in *Meneco*.

Reactions “R04568_C3_cytop” and “SO4tex”, were removed from the iDS372 and iJO1366 models respectively, to perform this assessment for growth validation. Meneco was then used to generate potential solutions for both models using two sets of “target” metabolites in parallel:

- The set of “target” metabolites comprised precursors and successors of the evaluated reaction in each model.
- The set of “target” metabolites was formulated based on the identification of dead-end metabolites by *BioISO*.

BiGG [14] universal database was used as the source of metabolic reactions for the test iJO1366 model, while KEGG [13] was used to solve gaps in the test iDS372 model.

## Availability and requirements

**Project name:** *BioISO*

**Project home page:** https://bioiso.bio.di.uminho.pt

**Operating system(s):** Platform independent

**Programming language:** Python

**Other requirements:** None

**License:** GNU GPL v3.0

**Any restrictions to use by non-academics:** None

The data generated or analysed during this study are included in this published article and its supplementary information files.

## List of abbreviations

BioISO: Biological networks constraint-based *In Silico* Optimization
GSMM: Genome-Scale Metabolic Model
FBA: Flux Balance Analysis
SBML: System Biology Markup Language
*wss*: total size of the whole search space
*ooss*: objective-oriented search space
*dem_ooss_*: ratio of dead-end metabolites for the objective-oriented search space
*dem_wss_*: ratio of dead-end metabolites for the whole-space search space
*br_ooss_*: ratio of blocked reactions for the objective-oriented search space
*br_wss_*: ratio of blocked reactions for the whole-space search space

## Declarations

### Competing interests

The authors declare that there is no conflict of interest.

### Funding

This study was supported by the Portuguese Foundation for Science and Technology (FCT) under the scope of the strategic funding of UIDB/04469/2020 unit and BioTecNorte operation (NORTE-01-0145-FEDER-000004) funded by the European Regional Development Fund under the scope of Norte2020 - Programa Operacional Regional do Norte. Fernando Cruz holds a doctoral fellowship (SFRH/BD/139198/2018) funded by the FCT.

## Acknowledgements

Not applicable

## Notes

### Competing Interest Statement

The authors have declared no competing interest.

