## Supplementary File 1 for "*BioISO*: an objective-oriented application for assisting the curation of genome-scale metabolic models"

### Supplementary file 1 – Comparing *BioISO* with state-of-the-art tools

#### Comparison with state-of-the-art tools

Comparing *BioISO* with state-of-the-art tools is challenging, as most algorithms aim to find and solve a wide range of different problems associated with the reconstruction of GSMMs. Most algorithms are commonly used in automatic gap-filling procedures. Hence, methods can be categorised according to their gap-finding and gap-filling methodology. Supplementary File S2 shows the main capabilities of several tools regarding the gap-finding and gap-filling methodologies, highlighting the differences between these applications.

Regarding the gap-finding methodology, methods can be classified according to the search type, search depth, and strategy to assert gaps. Guided- and exhaustive-searches are two distinct classes for the search type. Regarding the search depth, gap-finding methods can search for gaps in the whole metabolic network, or downstream and upstream of a user-defined reaction. Finally, four approaches are regularly used to assert whether a metabolite can be considered a gap:

- Stoichiometry: Stoichiometric analysis of the metabolite in the associated model reactions;
- Topological: Graph-based topological search for predecessors and successors;
- Consistency: Consistency between experimental data and phenotype simulations;
- Mixed: Graph-based topological search for predecessors and successors accounting for the metabolite’s stoichiometry in the associated reactions, together with FBA simulation of custom unbalanced reactions;

Although most tools can identify gaps in the metabolic network, *Smiley* [1] and *Mirage* [2] cannot be classified in terms of gap-finding. Both tools add new reactions to the model without an initial gap-scan, forcing model predictions to match the experimental data.

*BioISO*, *BiomassPrecursorCheck* [3], and *Meneco* [4] apply guided-search algorithms, as these tools regularly search for gaps or errors directly associated with a given objective/reaction. *BiomassPrecursorCheck* searches gaps in the reactions immediately downstream of the biomass reaction of a given model. Alternatively, *BioISO* seeks gaps downstream and upstream of a user-defined objective, allowing to almost search the whole metabolic network. *Meneco* searches for gaps according to a set of seed and target metabolites. However, the search depth may encompass the whole metabolic network. *gapFind/gapFill* [5], *fastGapFill* [6], and *Gauge* [7] are based on exhaustive searches. Thus, these methodologies identify gaps all over the metabolic network, regardless of a given objective.

*BioISO*, *biomassPrecursorCheck*, and *Meneco* check the metabolic network topological features to assert gaps. That is, these tools assert the existence of predecessors and successors of a given metabolite. Additionally, *BioISO* can identify both downstream and upstream metabolites from the user-defined objective by accounting for the metabolite’s stoichiometry in associated reactions. *BioISO* performs then multiple FBA’ simulations of custom unbalanced reactions during the topological search to evaluate whether a given metabolite is being consumed or produced. *gapFind/gapFill* and *fastGapfill* highlight gaps using a stoichiometry-like approach. These methods search the stoichiometric matrix for no-production and no-consumption metabolites. Alternatively, *Gauge* combines Flux Coupling Analysis and gene expression data to propose gaps in a draft GSMM.

The data requirements, free distribution, requirement for coding skills, and main output can be used to assess the usability of gap-finding tools. Only *BioISO* and *Meneco* are freely available, as the other methods rely on proprietary software, such as MATLAB (Mathworks®) or GAMS. Most of the gap-finding tools were implemented as Python packages, add-ons to the COBRA Toolbox [3], or GAMS files. It is worth noticing that *BioISO* is the only tool that does not require coding skills, as it is available as a webserver and in *merlin* software [8].

The main output of these tools consists of an array of missing metabolites and sets of potential solutions (often reactions retrieved from the metabolic data). *BioISO* is the only tool that provides a graphical user interface to visualise gaps and errors in a tree-based structure.

The gap-filling methods can be categorised according to the source and set of solutions to fulfil the metabolic network. The set of solutions offered by the gap-filling tools is often based on a minimal reaction set for a single gap. Alternatively, other methods consider a pan-metabolic network that assures flux through all metabolites. Thus, the solution set is often the result of two very different gap-filling approaches, namely the parsimonious and pruning approaches. These methods either add a reduced or large set of reactions to each gap, respectively. Regarding the latter approach, a pruning step is applied to reduce the large set of solutions. Whereas *Mirage* uses a pruning approach, most gap-filling methods use a parsimonious approach to complete a metabolic network gap.

All methods require a dataset of metabolic reactions to provide solution sets for fulfilling the draft metabolic network. Most methods consider the whole dataset of metabolic reactions retrieved usually from a biochemical database (e.g. KEGG [9], BiGG [10] or MetaCyc [11]). Other tools consider the reversible form of all reactions available in the metabolic model as an additional source of solutions. Besides a database of metabolic reactions, both *Gauge* and *Mirage* require gene expression data. In contrast, *Smiley* relies on growth phenotype data to identify minimal environmental conditions for which the model mispredicted growth and non-growth phenotypes.

It is worth noting that most tools addressed in this study have been designed explicitly for gap-filling procedures. These tools repeatedly warn users of the need to revise the set of candidate solutions for the numerous gaps and errors detected in the model. Alternatively, the main outcome can be a gapless metabolic model, not compliant with a high-quality bottom-up reconstruction.

Figure S1.1 exhibits these tools’ major strengths and weaknesses regarding the reconstruction of high-quality models by scientists without coding skills, using a bottom-up approach.

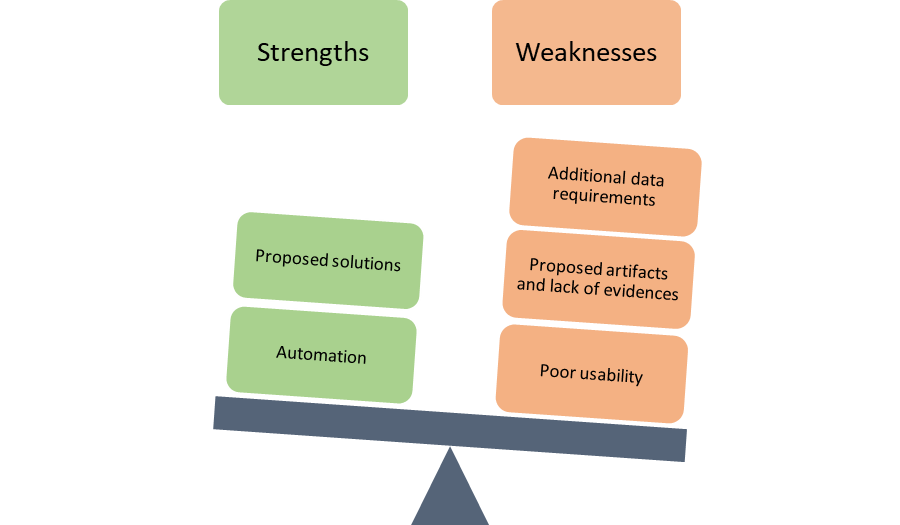

**Figure S1.1.** Strengths and weaknesses of gap-fill methods assessed in this work. Although the automatic proposal of solutions (set of reactions) for the numerous gaps identified by the gap-fill approaches might solve most issues, associated with the first steps of the GSMM reconstruction, the proposal of artefacts (namely, reactions and pathways without evidence) might raise other issues during model validation. Moreover, the requirement of additional data and coding skills, as well as the production of highly verbose and extensive outputs, hinders the usability of these tools for reconstructing high-quality models by wet-lab scientists.
