## Supplementary File 3 for "*BioISO*: an objective-oriented application for assisting the curation of genome-scale metabolic models"

### Supplementary file 3 – *BioISO*’s workflow, algorithm, and runtime analysis

The methodology for finding and assessing metabolites and reactions is detailed in the next sections. *BioISO*’s algorithm is described first together with samples of pseudo code that depict all tasks performed by the tool. Then, an analysis of *BioISO*’s relation-like algorithm’s complexity is described together with the recursion tree method visualisation.

#### Algorithm

The first algorithm labelled *BioISO* is the core logic supporting the methodology proposed in this work. A set of nodes, typically the metabolites associated with the reaction being evaluated, must be provided to initiate algorithm S3.1. The set of nodes is iterated one node at a time, and the reactions associated with each node are evaluated. In the next step, *BioISO* retrieves the set of reactants and products associated with each new reaction. As a result, a new node is created and assessed for each reactant and product. This workflow is repeated until reaching the stopping condition due to the recursive nature of the algorithm.

The stopping condition, *BioISO’s* depth parameter, represents the number of recursive calls performed by algorithm S3.1, when analysing the metabolic network. For instance, varying the algorithm’s depth from 1 to 3 allows running *BioISO* for shallow, guided or nearly exhaustive searches, depending on the metabolic network’s size and arborescence.

Algorithm S3.1: BioISO(nodes)

1: **Input:**

2:	*nodes*

4:

5: **for** each *node* in *nodes* **do**

6:	*nextNodes* $\leftarrow$  *Initialize EmptyArray()*

7:	*currentReactions* $\leftarrow$ *getNodeReactions*(*node*)

8:

9:	**for** each *reaction* in *currentReactions* **do**

10:

11:	*reactants* $\leftarrow$ *getReactants(reaction)*

*12:	products* $\leftarrow$ *getProducts(reaction)*

13:

14:	**for** each *reactant*, *product* in *reactants*, *products* **do**

15:	*nextNode* $\leftarrow$ *Initialize Node(reactant)*

*16:	nextNodeReactions* $\leftarrow$ *getReactions(nextNode, currentReactions)*

17:	*setNextReactions(nextNode, nextNodeReactions)*

18:	*nodeEval* $\leftarrow$ *fbaSimulReactant*(*nextNode*, *reactants*, *products)*

19:	*setEval*(*nextNode*, *nodeEval*)

20:	*setNextNode*(*node*, *nextNode*)

21:	*nextNodes.insert(nextNode)*

22:

23:	*nextNode* $\leftarrow$ *Initialize Node(product)*

24:	*nextNodeReactions* $\leftarrow$ *getReactions(nextNode, currentReactions)*

25:	*setNextReactions(nextNode, nextNodeReactions)*

26:	*nodeEval* $\leftarrow$ *fbaSimulProduct*(*nextNode*, *reactants*, *products)*

27:	*setEval*(*nextNode*, *nodeEval*)

28:	*setNextNode*(*node*, *nextNode*)

29:	*nextNodes.insert(nextNode)*

30:	**end for**

31:	**end for**

32:	*BioISO(nextNodes)*

33: **end for**

According to algorithm S3.2 (*testReactions*), the list of reactions to be retrieved next should not include the previous reactions, to avoid futile loops when creating the hierarchical structure. Furthermore, each reaction is assessed through a simple FBA simulation. The evaluation is positive if the reaction has flux; otherwise, *BioISO* considers that the reaction cannot attain the desired phenotype.

Algorithm S3.2: BioISO – testReactions (node, currentReactions)

1: **Input:**

2:	*node*

3:	*currentReactions*

4:

5: *reactions* $\leftarrow$  *findReactionsAssociatedWithNode*(*node*) #usually provided by a computational framework to analyze GSM models or biological networks

6: *nextReactionsList* $\leftarrow$  *Initialize EmptyArray()*

7:

8: **for** each *reaction* in *reactions* **do**

\# The next reactions list should contain the previous reactions list to prevent cyclic hierarchical structures

9:	**if** *reaction* not in *currentReactions* **then**

10:	*reactionEval* $\leftarrow$ *fbaSimul(reaction)*

*11:* 	*setEval*(*reaction*, *reactionEval*)

12:	add *reaction* to *nextReactionsList*

13:	**else**

14:	*continue*

15:	**end if**

16: **end for**

*BioISO* uses a more comprehensive approach to evaluate reactants (precursors) and products (successors), as demonstrated in algorithms S3.3 (*fbaSimulReactant*) and S3.4 (*fbaSimulProduct*), respectively.

A **precursor (reactant)** is considered as a positive assessment if the metabolic model can **produce** it (connected metabolite). Thus, a reactant is a product elsewhere in the metabolic network; otherwise, it would not be available for the objective reaction. Hence, *BioISO* creates an unbalanced reaction explicitly designed to allow the metabolite accumulation in the metabolic model.

The maximisation of this unbalanced reaction is the objective function of the FBA simulation. The model should synthesise the precursor metabolite, obtaining an optimal non-zero flux solution through the unbalanced reaction. In short, the evaluation is successful if the model can attain a positive flux in the surrogate reaction.

Furthermore, a similar reaction is included in the model for each reactant, to prevent the seldom cases in which all reactants are forcibly produced by reactions that produce/synthesise the assessed metabolite.

Likewise, *BioISO* creates unbalanced reactions that allow the uptake of all products associated with the evaluated reaction. These reactions are included in the model to cover up for the unlikely scenario that the model forcibly needs to consume/metabolise such products to synthesise the precursor.

For example, when evaluating reaction *R_11_*:

*R_11_*: I + J + L -> M	0 < *R_11_* < +∞

*BioISO* will evaluate the precursor *I* by adding an unbalanced reaction *R_13_*, which takes *I* as a reactant, and whose lower and upper bounds are set to zero and plus-infinity, respectively.

*R_13_*: I ->	0 < R_13_ < +∞

Since reaction *R_11_* has other reactants, similar reactions are added to the model, for each metabolite. Namely, *R_14_* and *R_15_* are added to the model as surrogate reactions for the accumulation of *J* and *L*, respectively. These metabolites can be produced only in the same reactions producing *I*. Similarly, *BioISO* will create an unbalanced reaction, namely *R_16_*, as surrogate reaction for the uptake of *M*. Again, to cover the unlikely chance that *M* is required to produce *I*.

*R_14_*: J ->	0 < R_14_ < +∞

*R_15_*: L ->	0 < R_15_ < +∞

*R_16_*: M ->	-∞ < R_16_ < 0

Finally, the objective function of the FBA consists of maximising reaction *R_13_*. Hence, a positive solution for this linear problem indicates a successful evaluation of metabolite *I* as a precursor when evaluating *R_11_*.

$$maximize \to R_{13}$$

Algorithm S3.3: BioISO – testReactant (node, reactants, products)

1: **Input:**

2:	*node*

3:	*reactants*

4:	*products*

5:

6: **for** *product* in *products* **do**

7:	*unbalancedReaction* $\leftarrow$ *createUnbalancedReaction(product)*

\# reaction formula: product <-

8:	*setBounds*(*unbalancedReaction*, -999999, 0)

9: **end for**

10: **for** *reactant* in *reactants* **do**

11:	**if** *reactant* is not *node* **then**

12:	*unbalancedReaction* $\leftarrow$ *createUnbalancedReaction(reactant)*

\# reaction formula: reactant ->

13: 	*setBounds*(*unbalancedReaction*, 0, 999999)

14:	**else**

15:	*continue*

16:	**end if**

17: **end for**

18:

19: *unbalancedReaction* $\leftarrow$ *createUnbalancedReaction(node)*

\# reaction formula: node ->

20: *setBounds*(*unbalancedReaction*, 0, 999999)

21: *reactantEval* $\leftarrow$ *fbaSimul(unbalancedReaction, maximize)*

On the other hand, a **successor (product)** is considered as a positive assessment if the metabolic model can **consume** it (connected metabolite). Thus, a product is a reactant elsewhere in the metabolic network. As described in the testing of precursor *I*, *BioISO* also creates an unbalanced reaction for the successor. However, this reaction is now explicitly designed to allow the metabolite **uptake** in the metabolic model. Thus, the minimisation of this uptake reaction is now the objective function of the FBA simulation. In other words, the model should now metabolise/consume the precursor metabolite, obtaining an optimal non-zero flux solution through the unbalanced reaction.

As described in the testing of precursor *I*, *BioISO* also creates unbalanced reactions that allow the uptake of all products as well as the accumulation of all reactants associated with the evaluated reaction.

For example, when evaluating reaction *R_11_*:

*R_11_*: I + J + L -> M;	0 < R_11_ < ∞

*BioISO* will evaluate first the successor *M* by adding the unbalanced reaction *R_13_*, which takes *M* as a reactant, and whose lower and upper bounds are set to minus-infinity and zero, respectively.

*R_13_*: M <-	-∞ < R_13_ < 0

As reaction *R_11_* does not have more products than metabolite *M*, only reaction *R_13_* is added to the model as an uptake reaction. *BioISO* adds next more unbalanced reactions that allow the accumulation of the reactants *I*, *J,* and *L*, namely *R_14_*, *R_15,_* and *R_16_*.

*R_14_*: I ->	0 < R_14_ < ∞

*R_15_*: J ->	0 < R_15_ < ∞

*R_16_*: L ->	0 < R_16_ < ∞

Finally, the objective function of the FBA simulation consists of minimising reaction *R_13_*. Hence, a negative flux solution for this linear problem indicates a successful evaluation of metabolite *M* as a successor (connected metabolite) when evaluating *R_11_*.

$$minimize \to R_{13}$$

*BioISO* implements a cache memory system of all simulations performed during the recursion. Thus, reactions and metabolites are only evaluated once for the specific role during the analysis.

Algorithm S3.4: BioISO – testProduct (node, reactants, products)

1: **Input:**

2:	*node*

3:	*reactants*

4:	*products*

5:

6: **for** *reactant* in *reactants* **do**

7:	*unbalancedReaction* $\leftarrow$ *createUnbalancedReaction(reactant)*

\# reaction formula: reactant ->

8:	*setBounds*(*unbalancedReaction*, 0, 999999)

9: **end for**

10: **for** *product* in *products* **do**

11:	**if** *product* is not *node* **then**

12:	*unbalancedReaction* $\leftarrow$ *createUnbalancedReaction(product)*

\# reaction formula: product <-

13: 	*setBounds*(*unbalancedReaction*, -999999, 0)

14:	**else**

15:	*continue*

16:	**end if**

17: **end for**

18:

19: *unbalancedReaction* $\leftarrow$ *createUnbalancedReaction(node)*

\# reaction formula: node <-

20: *setBounds*(*unbalancedReaction*, -999999, 0)

21: *productEval* $\leftarrow$ *fbaSimul(unbalancedReaction, minimize)*

#### BioISO runtime estimation

Finally, *BioISO* was implemented as a recursive relation-like computational method. Thus, it is possible to represent the algorithm by a runtime estimation function such as the one provided in notation 1 or visualized by applying the recursion tree method (Figure S3.1). Note that, the hierarchical tree-based structure presented in Figure S3.1 is based on the metabolic network depicted in Figure 1 of the manuscript.

In Figure S3.1, one can visualize that the amount of work done by *BioISO* at the first level is *n* and increases in the following levels. The even size of the tree, and each level’s cost divided by the sum of the amount of work per node per level, is also shown in Figure S3.1. Note that, the work of pooling the next precursors and successors is seemingly the same at each node.

Figure S3.1 also illustrates cases in which the function *BioISO* will not find precursors or reactants. As a result, such nodes are labelled as leaves or terminal nodes, and thus no further calls of algorithm S3.1 will be made.

The notation that describes all previous situations mathematically is represented as follows:

$T\left( n \right): 2T\left( n \right)-2\times l\times n+n$	(1)

Such that,

- $n$ is the amount of work
- $n$ is given by a function, *f(*$n$*)*
- $l$ is the number of leaves in the last levels

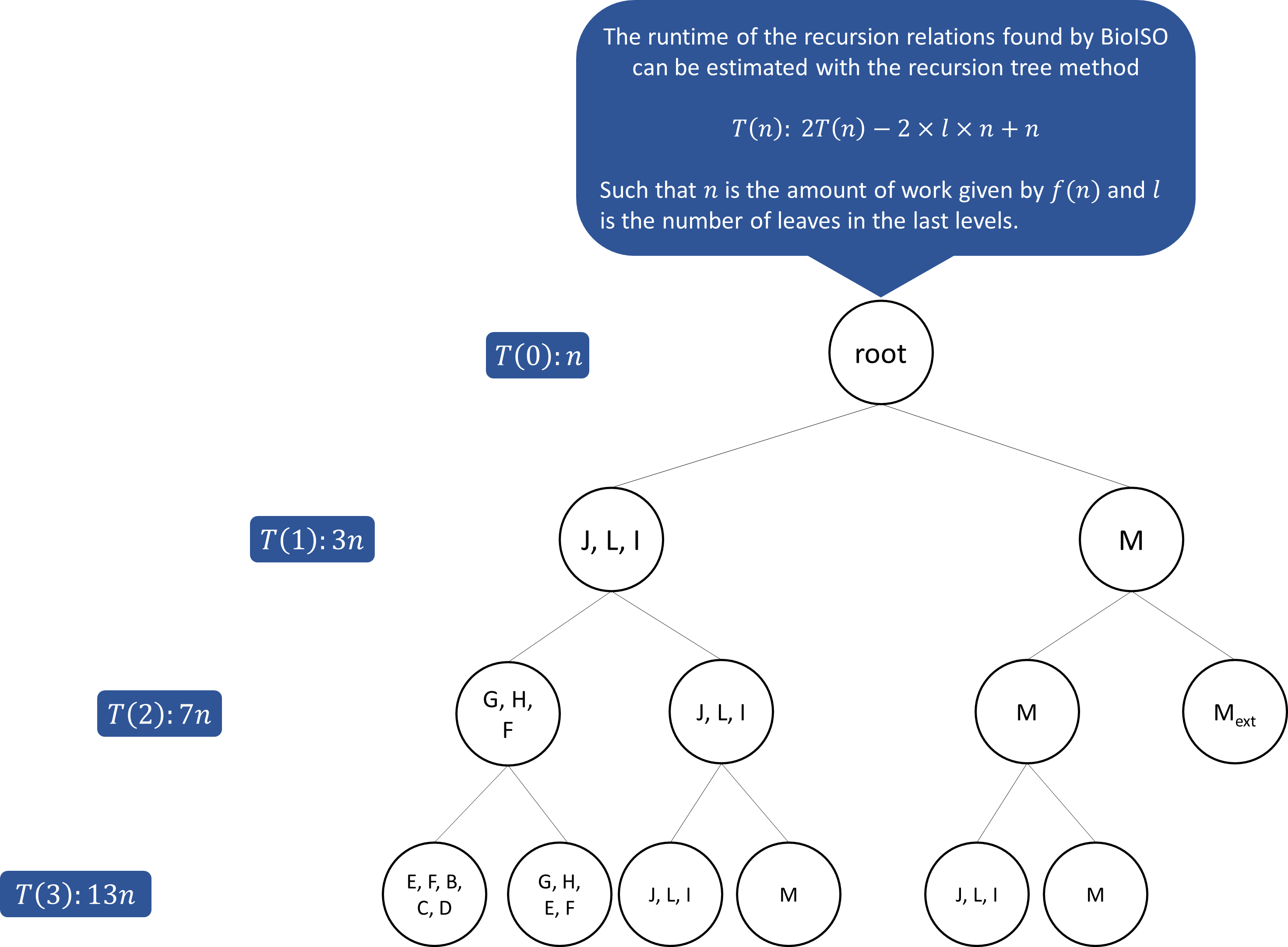

**Figure S3.1.** Overview of BioISO runtime estimation. The hierarchical tree-based structure imposed by the recursive relation-like computational method implemented in *BioISO* for the evaluation of reaction *R_11_*. Note that, the current hierarchical tree-based structure is based on the metabolic network depicted in Figure 1 of the manuscript. The recursion tree method is applied to visualize the distribution of a problem of size $n$ into multiple sub-problems (nodes) of similar size upon each recursion call. Each node in the recursion tree visualization stands for a sub-problem found by *f(*$n$*)* at each recursive call.
