## Supplementary File 4 for "*BioISO*: an objective-oriented application for assisting the curation of genome-scale metabolic models"

### Supplementary file 4 – Interpreting *BioISO* results

#### Running analysis in the available applications

GSM models derive from annotated genomes and biochemical data. As this information is often incomplete or misannotated, gaps and errors might appear in the metabolic network. Correspondingly, to obtain accurate and functional networks, these errors and inconsistencies must be thoroughly corrected. To the best of our knowledge, *BioISO* can accelerate and assist the gap-filling process in a user-friendly environment.

*BioISO* is currently implemented as a web-server and in the computational tool *merlin* [1].

#### Running analysis in the webserver

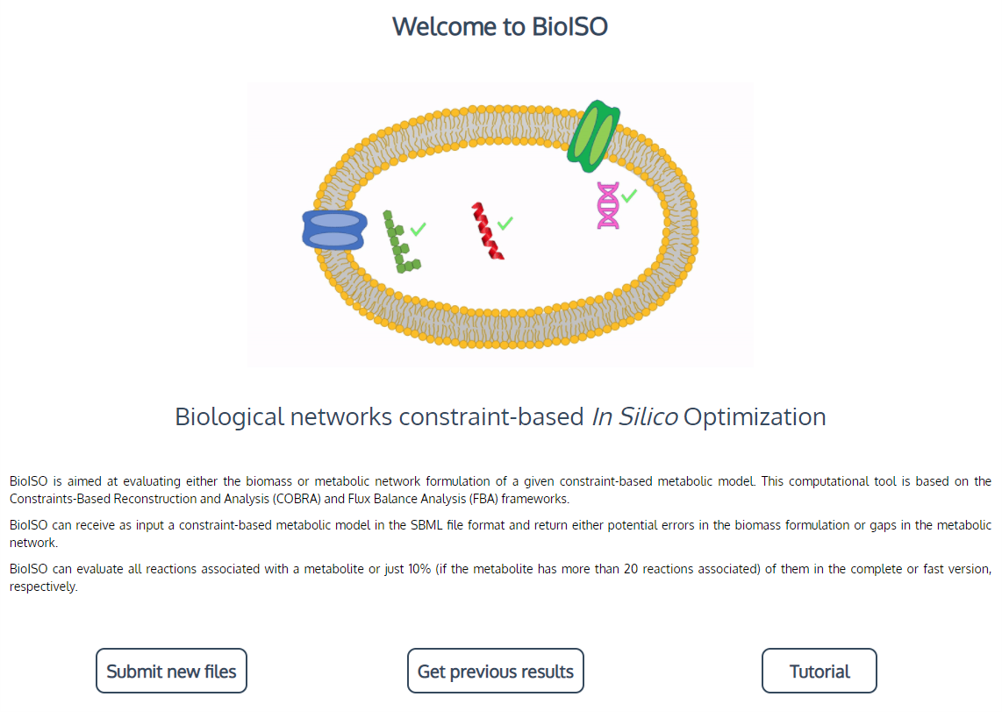

**Figure S4**.**1. -** Using *BioISO* webserver implementation - Homepage with *BioISO* logo and description. Two options are offered: “Submit new files” and “Get previous results”. The first option allows us to perform a new simulation, whereas the latter provides the results of previous simulations.

Step 1. Go to [https://BioISO.bio.di.uminho.pt/](https://bioiso.bio.di.uminho.pt/) on the web. A window such as the one in Figure S4.1 will be rendered.
Step 2. Press the “Submit new files” button to perform a new *BioISO* simulation.
Step 3. As shown in Figure S4.2, a set of parameters will be requested. Fill or choose the option in the boxes. Then, press the “Submit” button at the bottom.
Step 4. Wait for your results, this might take a few minutes. *BioISO* is running.
Step 5. Next, a view of the results will be rendered. As shown in Figures S4.3., S4.4., and S4.5., *BioISO* renders the information in interactions’ tables. The first table shows the metabolites associated with the optimized reaction. The “evaluation” field displays whether the metabolite is being produced (if it is reactant) or consumed (if it is a product). In short, *BioISO* highlights whether the metabolite is a dead-end metabolite or not.
Step 6. *BioISO* results. When the result of the optimized reaction is negative, as shown in Figure S4.6., its reactants and products evaluation table should be analysed. Negative evaluations in reactions indicate that these are not carrying flux (blocked reactions) and errors might be occurring. Negative evaluations in metabolites indicate that these are not being produced nor consumed (dead-end metabolites), depending on their role. (*) To get more detailed information about special cases in *BioISO* analysis, check the “Special Cases” section of the present document.
Step 7. Inspect the network considering the errors highlighted by *BioISO*.
Step 8. Run *BioISO* again to determine whether the network errors were corrected.
Step 9. Repeat this process until the results are similar to Figure S4.7.

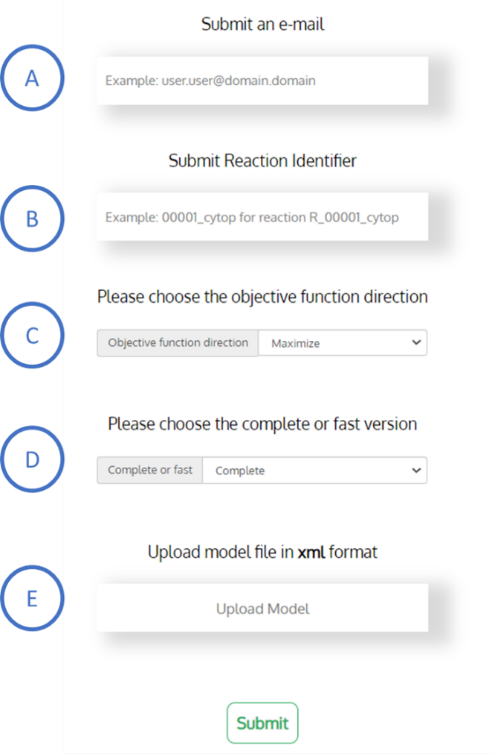

**Figure S4**.**2. -** Using *BioISO* webserver implementation. **A)** The user is requested to submit an e-mail. This e-mail will only be used to assign the results to a given user and not for registry purposes. **B)** The identifier of the reaction to be optimized is requested. *BioISO*’s analysis will focus on the optimization of this reaction to determine whether its reactants and products are being correctly produced and consumed, respectively. **C)** An objective function direction is required. The optimization can be either the maximization or the minimization of the objective function. **D)** *BioISO* offers two options: the complete and the fast search. The complete search analyses all the reactions associated with a given metabolite, while the fast search only evaluates 10 % of the reactions (if the metabolite has more than 20 reactions associated). **E)** The user is requested to submit a GSM model in the SBML format.

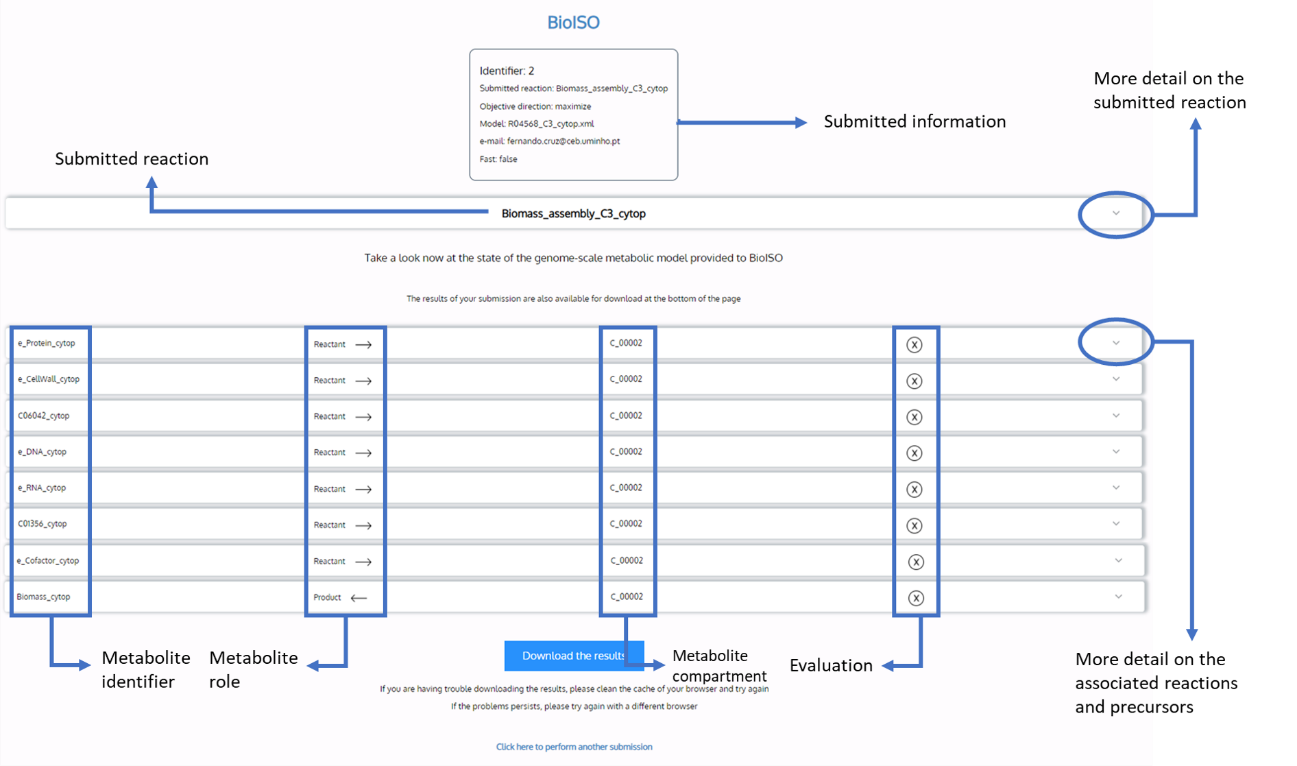

**Figure** S4.**3. -** Using *BioISO* webserver implementation. After the process is completed, a webpage opens with the simulation results. Here, the first level of results is rendered.

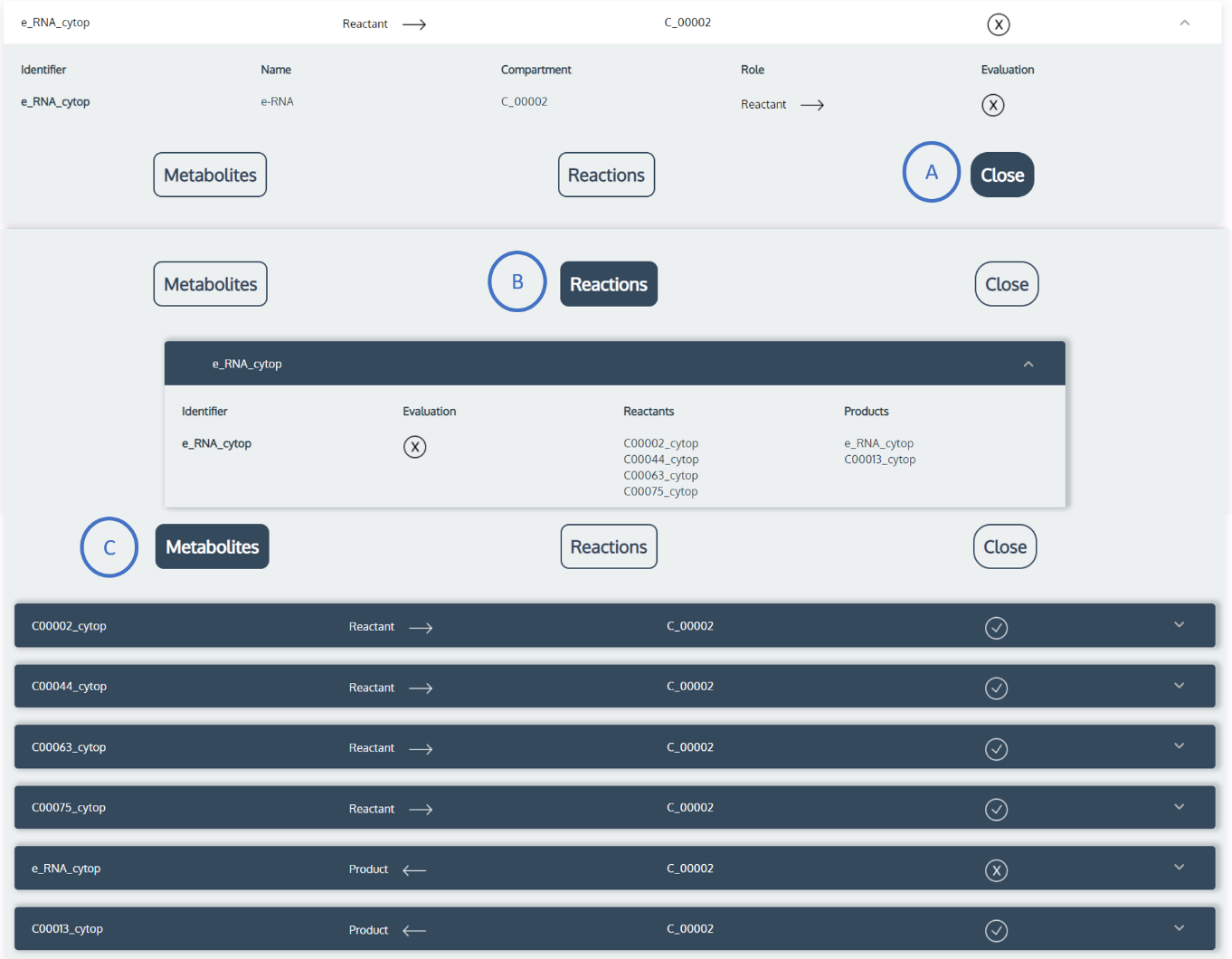

**Figure** S4.**4. –** Using *BioISO* webserver implementation. When the metabolite frame is expanded, buttons are disclosed indicating the type of information they can render. If the “Reactions” button is selected (**B**), the associated reactions and the respective evaluation is rendered, as well as the information about reactants and products. If the “Metabolites” button **C** is pressed, the second level of results is expanded, the reactions which produce this metabolite, and their evaluation will be rendered. Moreover, one can close these windows by pressing the button “Close” (**A**).

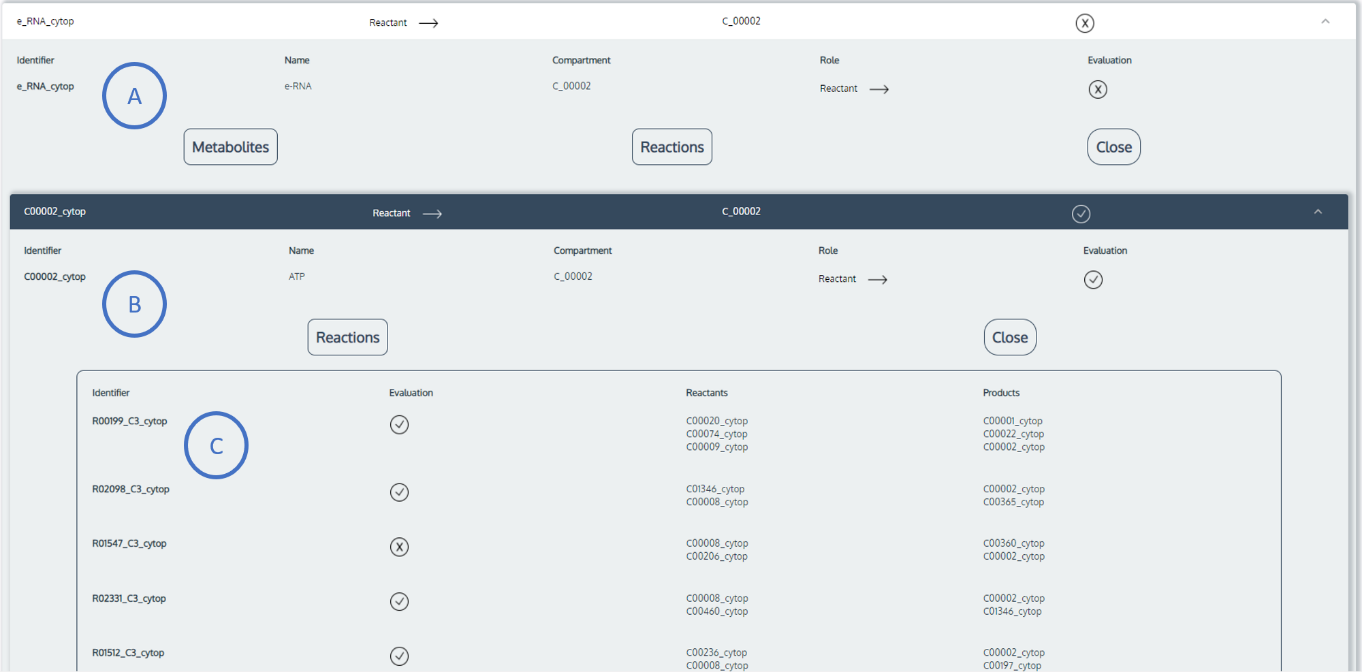

**Figure** S4.**5. –** Using *BioISO* webserver implementation. In this figure, the second level of results is shown. The metabolite being primarily inspected is shown in **A**, and the precursor in the window below (**B**). Here, information about their production or consumption is rendered. Lastly, a list of reactions producing precursor **B** are listed, as well as useful information about them (**C**).

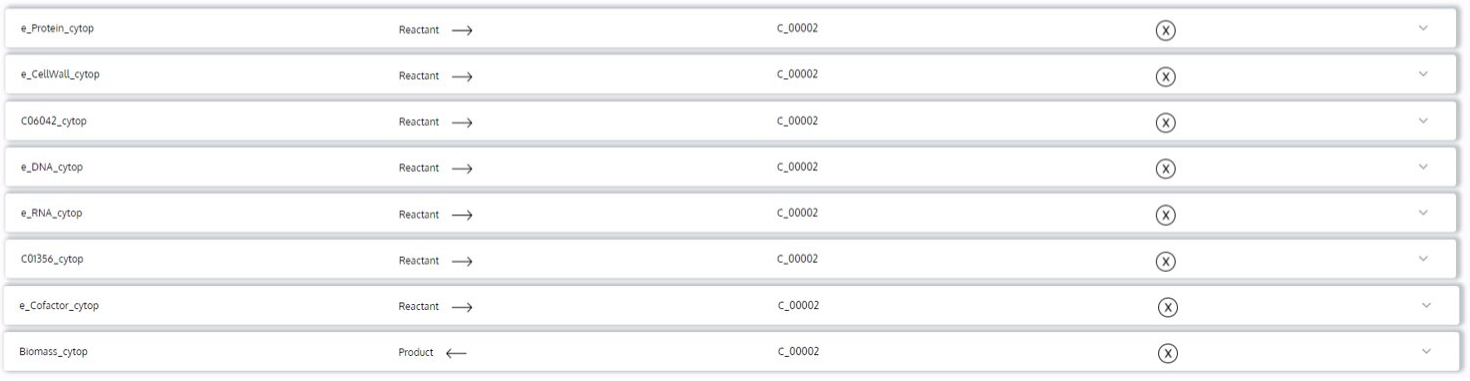

**Figure** S4.**6. –** Negative evaluation of metabolites (dead-end metabolites) - these metabolites are not being produced nor consumed (if the role is “Reactant”, their negative evaluation indicates that these are not being produced, on the other hand, if their role is “Product”, a negative evaluation indicates that these are not being consumed) – more information about the state of these metabolites can be checked expanding their window;

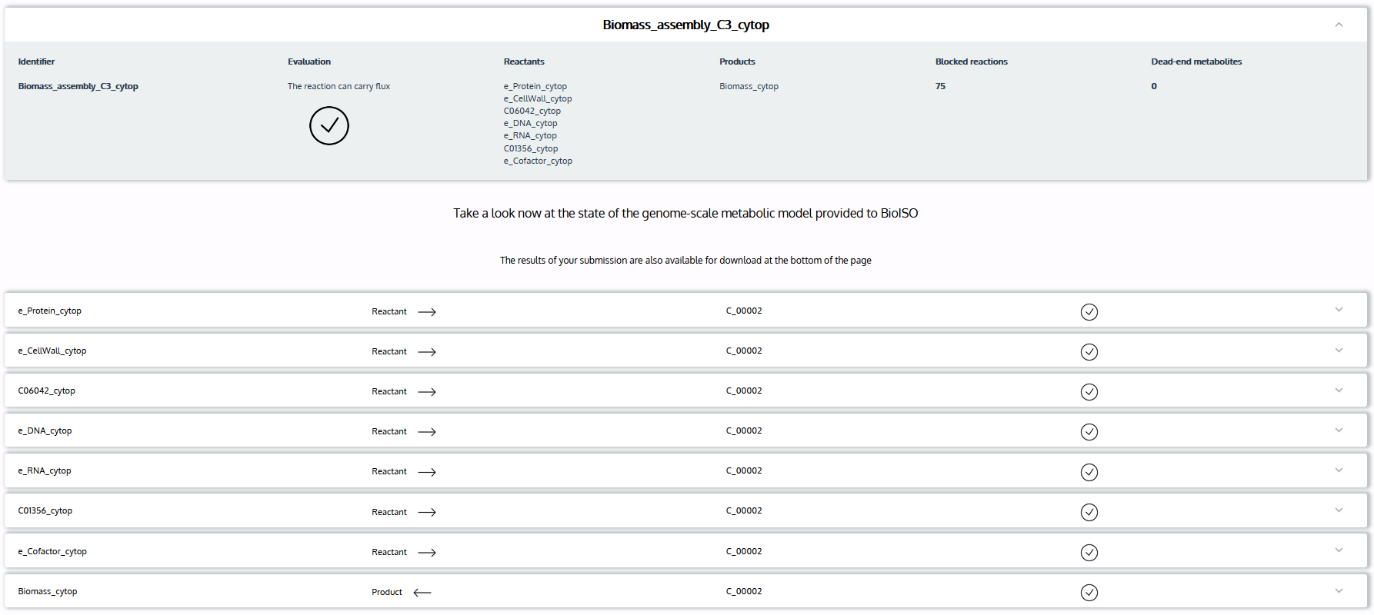

**Figure** S4.**7. –** *BioISO* simulation’s positive results. The simulated network has no errors nor gaps regarding biomass optimization.

#### Retrieve previous results from *BioISO* webserver

Step 1. Press the “Get previous results” button and two boxes will pop-up.
Step 2. Fill the first box with the submission identifier of the results you want to see.
Step 3. Write the e-mail used in the previous submission.
Step 4. Press “Submit” and a view with the results of the previous simulation will be rendered.

#### Running analysis in *merlin*

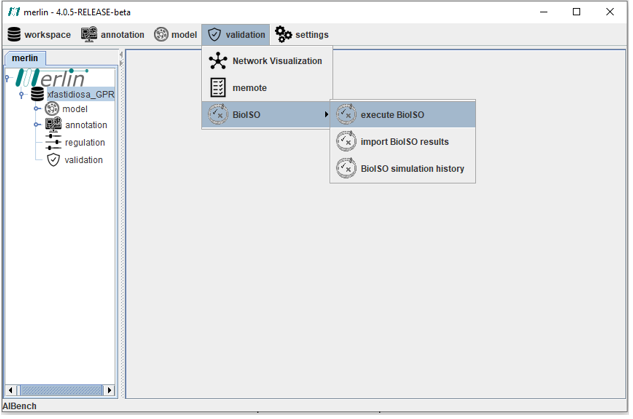

**Figure** S4.**8. -** Using *merlin’s* *BioISO* implementation. Selecting *BioISO* simulations in *merlin* graphical interface.

Step 1. Open the “Validation” menu in the top bar, then the *BioISO* menu will be available. Here, you will have the “execute *BioISO*” option. Select it. (Figure S4.8.)
Step 2. A menu will be opened where you can select the model, the reaction to be optimized, the objective (maximize or minimize), a checkbox to pick *BioISO*’s mode (fast or complete), and a text box to write a note about the simulation. (Figure S4.9)
Step 3. Wait for the results.
Step 4. *BioISO* results. The first level of results is rendered in *merlin’s* graphical interface. The first level of results is rendered in the main view (Figure S4.10.**A**), where, the metabolites evaluation is shown. More information about the reactions consuming or producing a given metabolite can be obtained by pressing the magnifier button. When pressed, a window is opened (Figure S4.10.**B**) with all the associated reactions and respective evaluations. More information about this reaction (Figure S4.10.**C**) can be obtained, once again, by pressing the magnifier button. An example of how to interpret *BioISO* results is described in Figure S4.11. (*) For more information about how to interpret *BioISO* results read the “Special Cases” section in the present document.
Step 5. Inspect your network considering the errors highlighted by *BioISO*.
Step 6. Run *BioISO* again to check whether your network errors were corrected.
Step 7. Repeat this process until your results look like the ones in Figure S4.10.**A.**

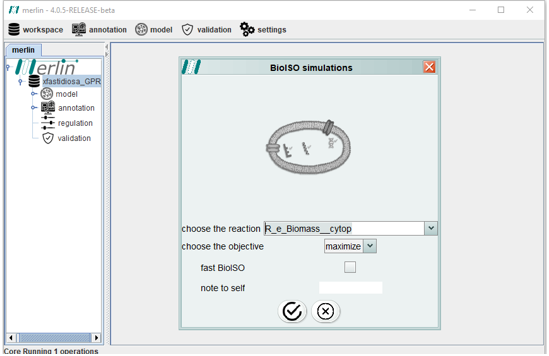

**Figure** S4.**9. –** Using *merlin’s* *BioISO* implementation. Selecting *BioISO* parameters in *merlin* graphical interface.

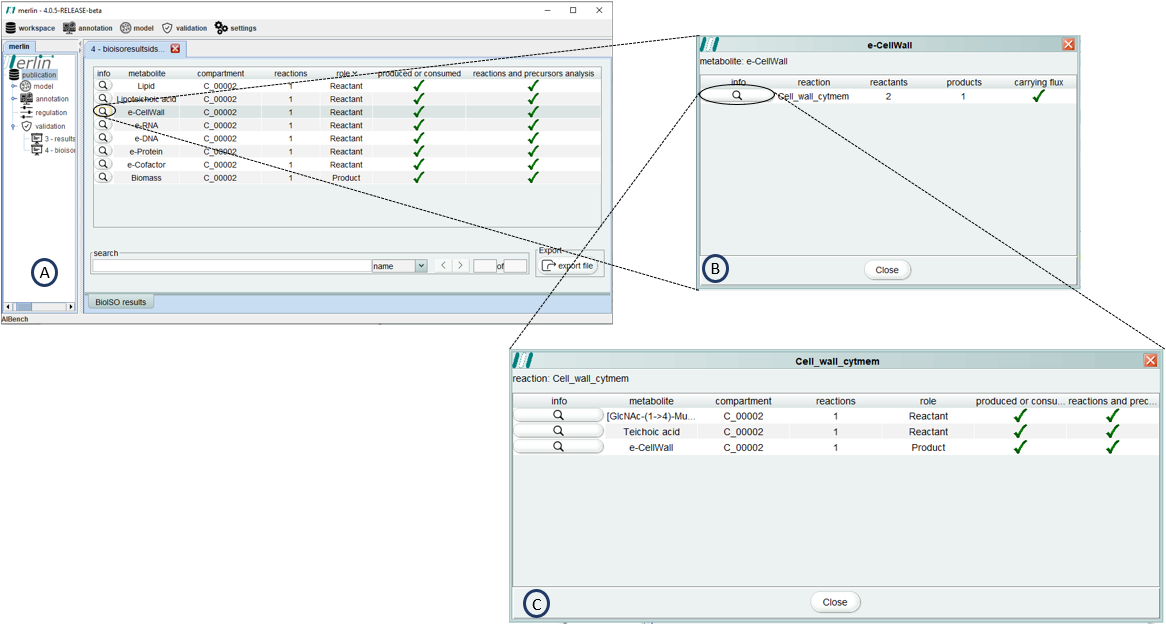

**Figure** S4.**10. –** Using *merlin’s* *BioISO* implementation. *BioISO* view in *merlin* graphical interface. Table **A** renders the evaluation of the metabolites (dead-end metabolites) associated with the optimized reaction. If the magnifier button is pressed, window **B**) is opened with the evaluation of the associated reactions (blocked reactions). Here, we can check whether the reactions associated with a specific metabolite are carrying flux (non-dead-end metabolite). Furthermore, more information about these reactions can be obtained by pressing the magnifier button (table **C**).

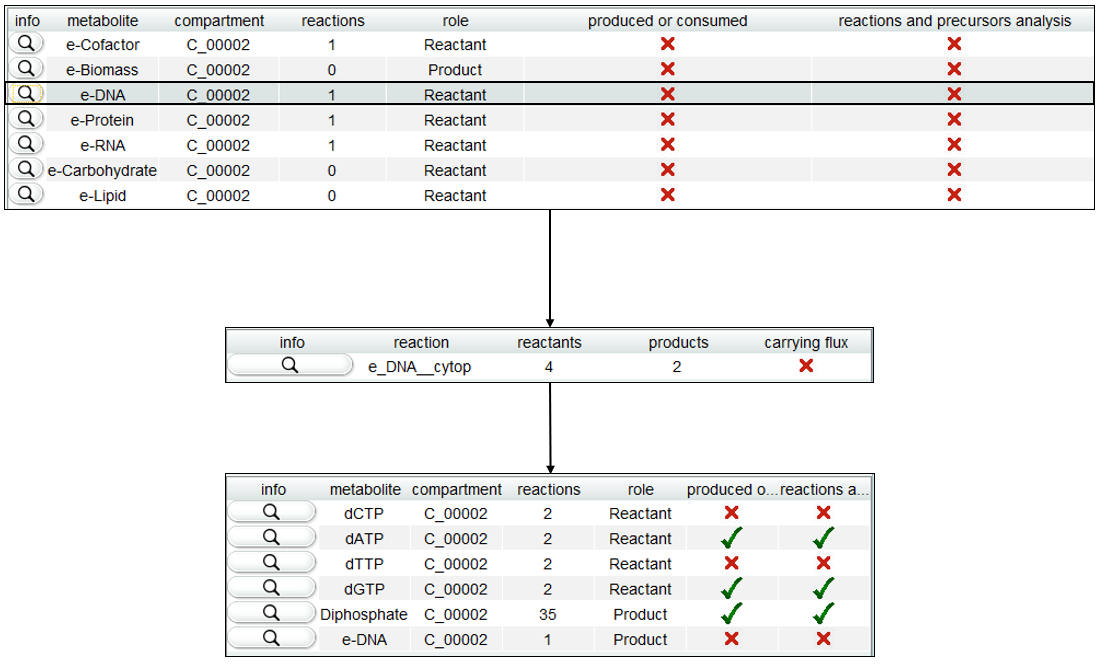

**Figure** S4.**11. –** Using *merlin’s* *BioISO* implementation. *BioISO*’s result example in *merlin.* This metabolic network does not produce any of the biomass precursors. For instance, if the magnifier button near the “e-DNA” metabolite is pressed, a reaction not carrying flux is shown. Correspondingly, by pressing the magnifier button in the new window, another table is rendered, identifying the reactants that are not being produced.

#### Special cases

There are special cases in the *BioISO* analysis that might impair the interpretation of the results. Thus, a detailed explanation will be hereby presented.

##### Case 1 – Metabolite with positive analysis and reactions with a negative evaluation

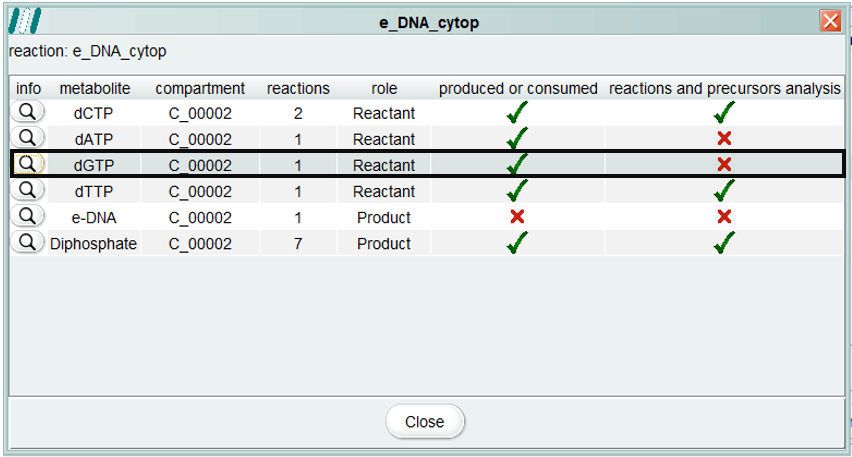

**Figure** S4.**12. –** Using *merlin’s* *BioISO* implementation. Example of case 1. Metabolite “dGTP” is being produced, however, there are errors in the precursors and reactions that are producing it.

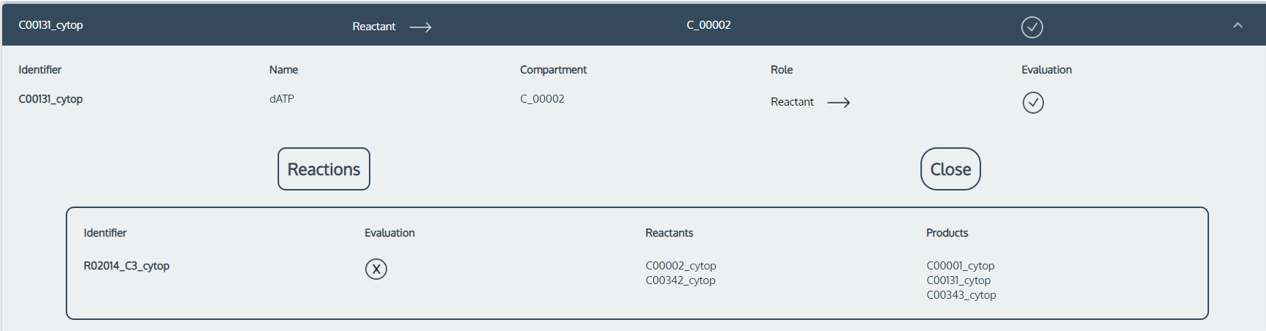

**Figure** S4.**13. –** Using *BioISO*’s web server implementation. Example of case 1. The metabolite “dATP” is being produced, however, there are errors in the precursors and reactions that are producing it.

This happens because reactions and metabolites are tested differently. Reactions are optimized towards a specific objective (maximization or minimization of flux), hence the presence of flux can be evaluated. In contrast, metabolites are tested individually within a given reaction. Moreover, note that it is likely that reactions in the model cannot attain any flux when optimized. However, the same reaction can obtain non-zero flux values when different objectives are used. This occurs because the metabolites involved in these reactions might be involved in cycles or cannot be accumulated or exported when optimizing such reaction.

As shown in Figure S4.13, the metabolite “dATP” is being correctly produced, however, the associated reaction (R02014_C3_cytop) cannot carry flux when optimized. On the other hand, when the objective function is to optimize the biomass reaction, R02014_C3_cytop can attain flux. Consequently, the metabolite evaluation is determined as positive because it is being produced towards the biomass reaction maximization. In this specific case, even though R02014_C3_cytop is not carrying flux when optimized, it certainly is when the objective function is the maximization of biomass production. The same logic can be applied to the example shown in Figure S4.12.

In short, the metabolite evaluation should be ultimately considered, as it is the most accurate evaluation. After correcting all the metabolites’ errors, go check again if their precursors are being correctly consumed and produced.

##### Case 2 – Metabolite with negative analysis and precursors with a positive analysis

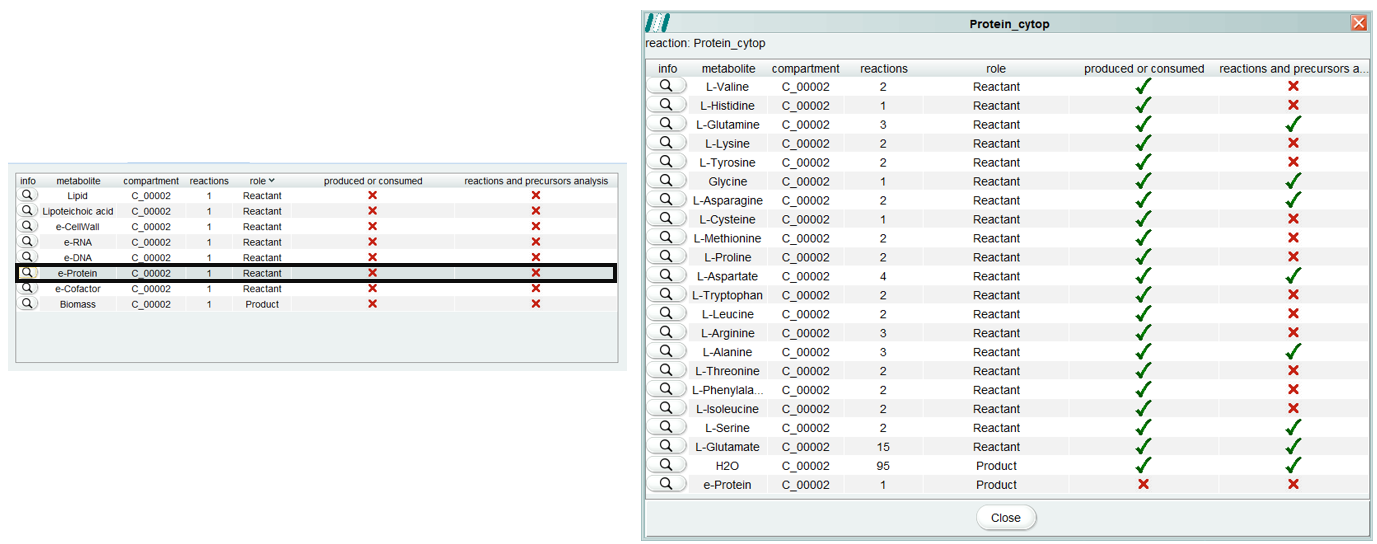

**Figure** S4.**14. –** Using *merlin* ‘s *BioISO* implementation. Example of case 2. Although the metabolite “e-Protein” has a negative evaluation, its precursors are all being produced and consumed correctly.

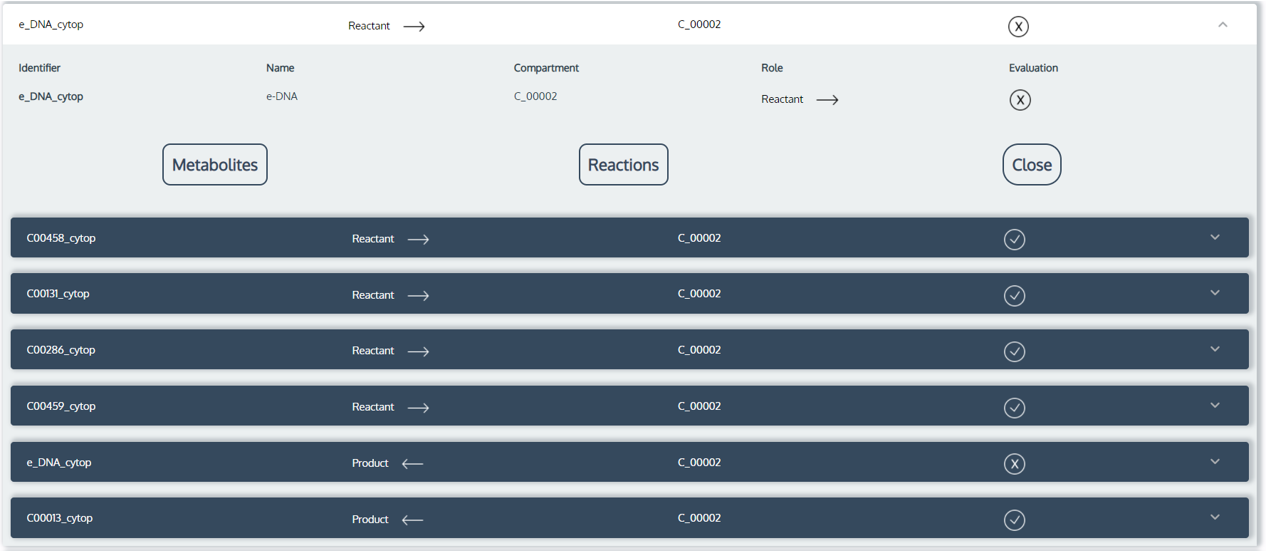

**Figure** S4.**15. –** Using *BioISO* web-server implementation. Example of case 2. Although the metabolite “e-DNA” has a negative evaluation, its precursors are all being produced and consumed correctly.

Whenever this case occurs, the metabolite evaluation is negative because the optimized reaction is not consuming the metabolite. Therefore, its reactants’ production and products’ consumption will solve the problem. In other words, the metabolite is being correctly produced, but it is not being consumed by any reaction.

As shown in Figure S4.14 and S4.15, a negative evaluation was assigned to the “e-Protein” and “e-DNA” metabolites, respectively. Regarding that their role is “product”, a negative evaluation indicates that they are not being consumed by any reaction. In the case presented in Figure S4.14, “e-Protein” is not being consumed by the biomass reaction, turning its evaluation negative, even though all its precursors are being correctly produced. The same logic can be applied to the example shown in Figure S4.15.
