## Supplementary File 5 for "*BioISO*: an objective-oriented application for assisting the curation of genome-scale metabolic models"

### Supplementary file 5 – *BioISO*’s validation methodology

### Genome-Scale Metabolic Models

Table S5.1 lists the Genome-Scale Metabolic Models (GSMMs) used for BioISO's validation. Each model's default environmental conditions were used to perform this validation. The carbon source was limited to 10 mmol _carbon source_/g/DW when not defined in the environmental conditions.

Table S5.1. Genome-Scale Metabolic Models (GSMMs) used during BioISO validation.

| **Model** | **Organism** | **Year** | **Reference** | **Environmental Conditions** |
| --- | --- | --- | --- | --- |
| iDS372 | *Streptococcus pneumoniae* | 2019 | [1] | Wild-type anaerobiosis defined in the published model |
| iJO1366 | *Escherichia coli* | 2011 | [2] | Wild-type anaerobiosis defined in the published model |
| iBsu1103 | *Bacillus subtilis* | 2009 | [3] | -10 mmol _nutrient source_/g/DW for all nutrients; |
| iTO977 | *Saccharomyces cerevisiae* | 2013 | [4] | Wild-type aerobiosis defined in the published model (-10 mmol_glucose_/g/DW) |
| iOD907 | *Kluyveromyces lactis* | 2014 | [5] | Wild-type aerobiosis (-10 mmol_glucose_/g/DW); |

### Computational software

BioISO was compared with state-of-the-art gap-finding and gap-filling tools. Table S5.2 lists the tools selected for comparison, which have been categorised into exhaustive-search and guided-search, according to the supplementary material S3. The latest version of each tool, together with IBM's CPLEX solver (version 12.10.0), on an eight-core Intel i7-9700 processor, was used in this assessment.

Table S5.2. State-of-the-art gap-finding and gap-filling tools used in BioISO's validation procedure.

| Tool | Search | Distribution | Year | Reference |
| --- | --- | --- | --- | --- |
| *Meneco* | Guided-search | Python package version 2.0.0 | 2017 | [6] |
| *fastGapFill* | Exhaustive-search | fastGapFill’s COBRA Toolbox version 3.0.0 | 2014 | [7] |

### Analytical methodology

#### Objective functions

Tables S5.3 and S5.4 exhibit the identifiers of the evaluated reactions for each model. Two main objective functions were assessed namely growth maximisation and compound production maximisation.

The wild-type growth rate, as well as compound production rates, were obtained after simulating the biomass and compound exchange reactions, respectively.

Table S5.3. Reactions evaluated when analysing growth maximisation with BioISO.

| Model | Reaction evaluated | Growth rate (h^-1^) |
| --- | --- | --- |
| iDS372 | Biomass_assembly_C3_cytop | 1.2095 |
| iJO1366 | Ec_biomass_iJO1366_core_53p95M | 0.2415 |
| iBsu1103 | bio00006 | 15.2068 |
| iTO977 | CBIOMASS | 0.9969 |
| iOD907 | Biomass_cyto | 1.0128 |

Table S5.4. Reactions evaluated when analysing compound production maximisation with BioISO.

| Model | Compound of interest | Compound exchange reaction | Flux rate (mmol/gDW/h) | Reaction evaluated |
| --- | --- | --- | --- | --- |
| iDS372 | Lactate | EX_C00186_extr | 66.4018 | R00703_C3_cytop |
| iJO1366 | Acetate | EX_ac_LPAREN_e_RPAREN_ | 26.7231 | ACKr |
| iBsu1103 | Acetate | EX_cpd00029_e | 1000.0 | rxn00227 |
| iTO977 | CO2 | CO2xtO | 60.0 | PDA1_2 |
| iOD907 | Citrate | EX_C00158_extr | 6.6667 | R00351_C3_cyto |

#### Creating models with gaps

The following methodology was used to create incomplete models for the tools' assessment:

1. A set of essential reactions was retrieved for each "model/objective function" pair using COBRApy's built-in method (*single_reaction_deletion*). Seldom, several essential reactions had to be removed from the model, as a single essential reaction for the objective function was not found. Table S5.5 highlights the additional reactions removed from each model.
2. Five reactions were randomly selected from the pool of essential reactions. When the pool size was less than five, all essential reactions were selected.
3. Each reaction was removed from the model, creating five incomplete replicas of the original model.

Table S5.5. Additional reactions were removed from each model to obtain a pool of essential reactions for the compound production objective function.

| **Model** | **Organism** | **Deleted reactions** |
| --- | --- | --- |
| iBsu1103 | *Bacillus subtilis* | rxn00152  rxn00173 |
| iOD907 | *Kluyveromyces lactis* | R00355_C3_cyto  R00344_C3_cyto  T01268_C4_mito  R01731_C3_cyto |

#### Evaluation metrics

Two metrics were proposed to evaluate the gap-finding tools’ performance, namely the ratio of dead-end metabolites and the ratio of blocked reactions. These metrics were used to quantify the search space associated with model debugging of gaps and errors.

The total size of the whole search space (*wss*) represents the number of metabolites or reactions available in the model. Whereas the objective-oriented search space (*ooss*) represents the number of metabolites or reactions covered by a guided-search tool during its objective-oriented search. As shown in equation 1), the ratio of dead-end metabolites for the objective-oriented search space (*dem_ooss_*) is a function of the number of metabolites that a guided-search tool evaluates as unsuccessful (dead-end metabolite) divided by the size of the *ooss*. The ratio of dead-end metabolites for the whole search space (*dem_wss_*) is a function of the number of found dead-end metabolites divided by the *wss*, as described in equation 2).

A low ratio of dead-end metabolites is associated with a small effort to find the error in the metabolic network. Thus, gap-finding and gap-filling tools should get low ratios to be compliant with high-quality bottom-up GSMMs reconstructions that often involve minor modifications.

Equations 3) and 4) describe a similar approach to calculate the ratio of blocked reactions for the objective-oriented search (*br_ooss_*) and the whole search (*br_wss_*) spaces, respectively.

As fastGapFill does not perform an objective-oriented search, the scores for objective-oriented searches (equations 1 and 3), are identical to whole searches’ scores.

${dem}_{ooss}= \frac{\sum DeadEndMetabolites}{\sum CoveredMetabolies}$	1)

${dem}_{wss}= \frac{\sum DeadEndMetabolites}{\sum Metabolies}$	2)

${br}_{ooss}= \frac{\sum BlockedReactions}{\sum CoveredReactions}$	3)

${br}_{wss}= \frac{\sum BlockedReactions}{\sum Reactions}$	4)
